# Dissecting embryonic and extra-embryonic lineage crosstalk with stem cell co-culture

**DOI:** 10.1101/2023.03.07.531525

**Authors:** Yulei Wei, E Zhang, Leqian Yu, Baiquan Ci, Lei Guo, Masahiro Sakurai, Shino Takii, Jian Liu, Daniel A. Schmitz, Yi Ding, Linfeng Zhan, Canbin Zheng, Hai-Xi Sun, Lin Xu, Daiji Okamura, Weizhi Ji, Tao Tan, Jun Wu

## Abstract

Faithful embryogenesis requires precise coordination between embryonic and extraembryonic tissues. Although stem cells from embryonic and extraembryonic origins have been generated for several mammalian species(Bogliotti et al., 2018; Choi et al., 2019; Cui et al., 2019; Evans and Kaufman, 1981; Kunath et al., 2005; Li et al., 2008; Martin, 1981; Okae et al., 2018; Tanaka et al., 1998; Thomson et al., 1998; Vandevoort et al., 2007; Vilarino et al., 2020; Yu et al., 2021b; Zhong et al., 2018), they are grown in different culture conditions with diverse media composition, which makes it difficult to study cross-lineage communication. Here, by using the same culture condition that activates FGF, TGF-β and WNT signaling pathways, we derived stable embryonic stem cells (ESCs), extraembryonic endoderm stem cells (XENs) and trophoblast stem cells (TSCs) from all three founding tissues of mouse and cynomolgus monkey blastocysts. This allowed us to establish embryonic and extraembryonic stem cell co-cultures to dissect lineage crosstalk during early mammalian development. Co-cultures of ESCs and XENs uncovered a conserved and previously unrecognized growth inhibition of pluripotent cells by extraembryonic endoderm cells, which is in part mediated through extracellular matrix signaling. Our study unveils a more universal state of stem cell self-renewal stabilized by activation, as opposed to inhibition, of developmental signaling pathways. The embryonic and extraembryonic stem cell co-culture strategy developed here will open new avenues for creating more faithful embryo models and developing more developmentally relevant differentiation protocols.

## INTRODUCTION

In mammals, embryogenesis is accompanied by the establishment and loss of pluripotency, a transient property enabling embryonic epiblast cells to generate all cells in an adult organism. Pluripotent epiblast cells can be propogated in vitro in a spectrum of pluripotent stem cell (PSC) states with different culture conditions (Pera and Rossant, 2021; Wu and Izpisua Belmonte, 2015). PSCs with unique molecular and functional features provide us with invaluable in vitro models to study early mammalian development with great potential for regenerative medicine.

Despite the potential, however, PSCs exist in the artificial milieu of cell culture, which is vastly different from the *in vivo* environment. One notable difference is the lack of communication with extraembryonic cells in cultured PSCs. Extraembryonic tissues play crucial roles in embryo patterning and size control during early embryogenesis (Rivera-Pérez and Hadjantonakis, 2015; Shahbazi and Zernicka-Goetz, 2018). Consequently, PSC-only models suffer from several culture artifacts that potentially limit their potential, e.g., unsynchronized and disorganized differentiation, and unfettered growth.

To overcome these limitations, for the first time, we succeeded in the derivation of stem cells from embryonic and extraembryonic tissues using the same condition from both mouse and cynomolgus monkey blastocysts. This enabled the culture of embryonic and extraembryonic stem cells in the same environment, and thereby opening the door for dissecting their direct communication.

## RESULTS

### A common culture for all blastocyst lineages

We previously reported *de novo* derivation of intermediate embryonic stem cells (ESCs) from mouse blastocysts by activating <u>F</u>GF, <u>T</u>GF-β and <u>W</u>NT pathways (FTW-mESCs)(Yu *et al*., 2021b). To enrich the FTW-mESC population, a MEK inhibitor (PD0325901) was added to suppress the proliferation of extra-embryonic cells(Yu *et al*., 2021b). Withdrawal of PD0325901 during FTW-mESCs derivation resulted in the co-appearance of epiblast-like cells (ELCs), trophoblast-like cells (TLCs), and extraembryonic endoderm-like cells (XLCs) in the blastocyst outgrowth (Figure S1A). After manual picking, separate cultivation, and further passaging in the FTW condition, stable e<u>x</u>traembryonic <u>en</u>doderm stem cells (designated as FTW-mXENs), trophoblast <u>s</u>tem <u>c</u>ells (designated as FTW-mTSCs), and FTW-mESCs could be derived from a single mouse blastocyst (Figure 1A and 1B). FTW-mXENs, FTW-mTSCs and FTW-mESCs expressed extraembryonic endoderm, trophoblast, and epiblast markers, respectively (Figure 1C and Figure S1B to S1E). Next, we compared FTW-mXENs and FTW-mTSCs with mXENs and mTSCs in conventional conditions(Chiu et al., 2010; Niakan et al., 2013) (Figure S1F to S1I). FTW-mTSCs and FTW-mXENs expressed their respective lineage markers at comparable levels to mTSCs and mXENs, respectively (Figure S1J). Although similar proliferation rates were observed for mTSCs and FTW-mTSCs, FTW-mXENs exhibited a significantly shorter doubling time than mXENs (Figure S1K). After random differentiation of FTW-mXENs *in vitro,* several parietal endoderm (PE) and visceral endoderm (VE) related genes were upregulated(Artus et al., 2012) (Figure S1L). Following subcutaneous injection into an immunodeficient NOD-SCID mouse, FTW-mXENs formed a teratoma-like tissue mass (designated as XEN-teratoma) after 3 months (Figure S1M). Immunofluorescence (IF) analysis revealed that within the XEN-teratoma many cells expressed alpha-fetoprotein (AFP)(Kwon et al., 2006), some cells were GATA6+, and others stained positive for yolk sac markers FOXA1 and/or COL6A1 (Figure S1N and S1O). We did not detect T+ (mesoderm) or PAX6+ (ectoderm) cells in the XEN-teratoma (Figure S1M). For FTW-mTSCs, after random differentiation *in vitro*, multinucleated cells appeared (Figure S1P) and genes related to trophoblast differentiation were upregulated(Cui *et al*., 2019) (Figure S1R). FTW-mTSCs also formed a tissue mass (TSC-teratoma) containing multinucleated trophoblast giant-cell like cells two weeks after injection into a NOD-SCID mouse (Figure S1Q). Next, we performed blastocyst injections and found that GFP (green fluorescent protein) or mKO (monomeric Kusabira-Orange) labeled FTW-mXENs and FTW-mTSCs contributed to chimera formation in the yolk sac and placenta tissues of mouse conceptuses (E7.5 and E11.5), respectively (Figure 1D to 1G, Figure S1S and Table S1).

**Figure 1.**
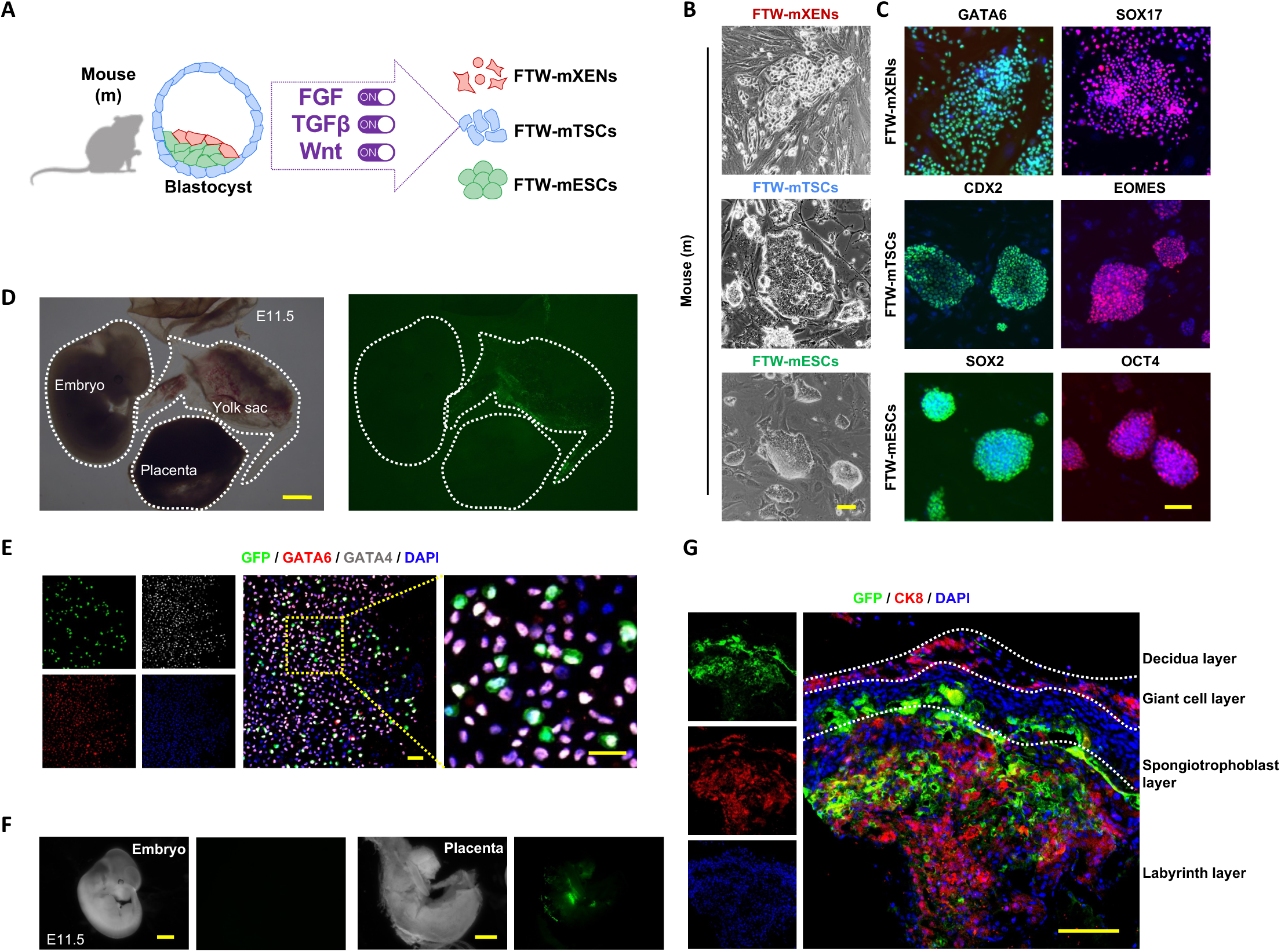
Derivation and characterization of embryonic and extraembryonic stem cells from mouse blastocyst. (A) Schematic of derivation of FTW embryonic and extraembryonic stem cell lines from mouse blastocyst. (B) Representative bright field (BF) images of colonies of FTW-mXENs, FTW-mTSCs and FTW-mESCs. Scale bar, 100 µm. (C) Representative immunofluorescence (IF) images of extraembryonic endoderm (GATA6 and SOX17), trophoblast (EOMES and CDX2), and epiblast (SOX2 and OCT4) lineage markers in FTW-mXENs (top), FTW-mTSCs (middle), and FTW-mESCs (bottom), respectively. Scale bar, 100 µm. (D) and (F) Representative BF and fluorescence images showing chimera contribution of GFP-labeled FTW-mXENs (D) and FTW-mTSCs (F) to E11.5 mouse conceptuses. Scale bar, 1 mm. (E) IF staining of a chimeric yolk sac membrane for GFP, GATA6 and GATA4. (G) IF staining of a chimeric sagittal placental section for CK8 and GFP. Different layers of the placenta are delineated by dotted lines. Scale bar, 100 µm. See also Figure S1.

In sum, we discovered a common culture condition could support *de novo* derivation and long-term culture of embryonic and extraembryonic stem cells from mouse blastocysts.

### Transcriptome profiling

We next performed bulk RNA-sequencing (RNA-seq) to examine the global transcriptional profiles of FTW stem cells and compared them with published datasets from established mouse embryonic and extraembryonic stem cells(Anderson et al., 2017; Bao et al., 2018; Cruz-Molina et al., 2017; Cui *et al*., 2019; Kubaczka et al., 2015; Wu et al., 2015; Wu et al., 2011; Ye et al., 2018; Zhao et al., 2015; Zhong *et al*., 2018). We found that FTW-mXENs, FTW-mTSCs and FTW-mESCs expressed their respective lineage markers and clustered together with stem cells derived from the same tissue of origin (Figure 2A and Figure S2A, S2B). Based on comparison with *in vivo* datasets, we found FTW-mESCs, FTW-mTSCs and FTW-mXENs were transcriptionally most closely related to E5.5 epiblast (EPI), E5.25-E5.5 extraembryonic ectoderm (ExE)(Cheng et al., 2019) and E5.5 VE(Mohammed et al., 2017), respectively (Figure S2C and S2D). To resolve the transcriptional states of FTW stem cells, we performed single-cell RNA-sequencing (scRNA-seq) and compared with published single-cell transcriptomes derived from E3.5-E6.5 mouse embryos(Nowotschin et al., 2019). Consistent with bulk RNA-seq results, we found that FTW-mESCs, FTW-mTSCs and FTW-mXENs showed the highest correlation with E5.5 EPI, E5.5 TE (trophectoderm) and E5.5 PrE (primitive endoderm), respectively (Figure 2B). Uniform manifold approximation and projection (UMAP) analysis further revealed FTW stem cells clustered closely with cells from their respective lineages in E5.5 mouse conceptuses (Figure S2E). To examine the temporal steps of FTW stem cells derivation, in addition to established FTW stem cell lines (passage 10), we performed scRNA-seq of blastocysts (day 0) and blastocyst outgrowths (day 8). Our analysis revealed clear segregation of ELCs, TLCs and ELCs in blastocyst outgrowths (Figure 2C). Slingshot(Street et al., 2018) pseudotime analysis was performed to delineate differentiation trajectories during derivation. Interestingly, in blastocyst outgrowths, ELCs have already acquired similar transcriptional states of stable FTW-mESCs while TLCs and ELCs clustered separately from established FTW-mTSCs and FTW-mXENs, respectively (Figure 2C). Next, we identified stage specific genes and enriched Gene Ontology (GO) terms. Notably, several common stage-specific features across all lineages emerged in our analysis, e.g. blastocysts were enriched with terms related to modification of DNA, RNA and/or protein, suggesting active gene transcription and translation, and dynamic signaling activities in these cells; cells in blastocyst outgrowths shared terms related to actin dynamics, which implicates active cell movement and dynamic changes in cell shape; established FTW stem cells, on the other hand, are characterized by terms related to glycolysis and hypoxia, which is indicative of their stabilization in the FTW condition (Figure 2D to 2F).

**Figure 2.**
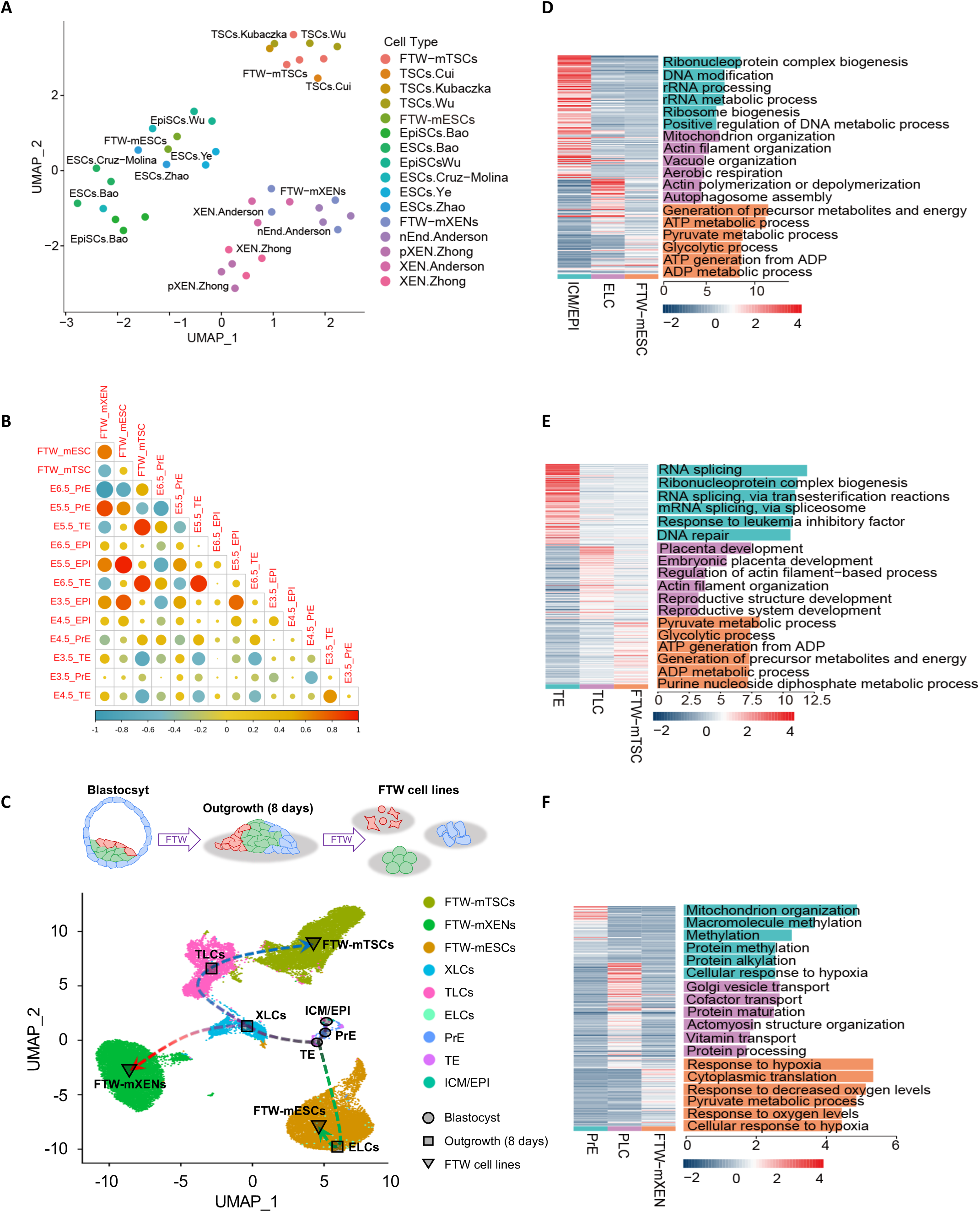
Transcriptomic analyses of mouse FTW stem cells. **(A)** UMAP visualization of integrative analysis of FTW-mXENs, FTW-mTSCs, FTW-mESCs and published datasets of mPSCs(Bao *et al*., 2018; Cruz-Molina *et al*., 2017; Wu *et al*., 2015; Ye *et al*., 2018; Zhao *et al*., 2015), mTSCs(Cui *et al*., 2019; Kubaczka *et al*., 2015; Wu *et al*., 2011) and mXENs(Anderson *et al*., 2017; Zhong *et al*., 2018). (B) Correlation analysis of FTW-mXENs, FTW-mTSCs and FTW-mXENs with published scRNA-seq dataset of in vivo mouse embryos from different stages. (C) UMAP analysis of mouse blastocysts, blastocyst outgrowths, and stable (passage 10) FTW-mESCs, FTW-mXENs and FTW-mTSCs. (D) to (F) Heatmaps and GO terms analysis of stage-specific (blastocyst, blastocyst outgrowth and stable FTW stem cells) genes from three lineages during FTW stem cells derivation. See also Figure S2.

Taken together, these transcriptomic analyses confirmed lineage identities, revealed temporal properties and derivation dynamics of FTW-mESCs, FTW-mTSCs and FTW-mXENs.

### Cross-lineage stem cell co-cultures

Having FTW-mESCs, FTW-mXENs and FTW-mTSCs derived and maintained in the same condition enabled us to establish co-cultures to study intercellular communications between embryonic and extraembryonic lineages (Figure 3A). To this end, we labeled FTW-mESCs with GFP and subjected them to co-culture with mKO-labeled FTW-mXENs, in the presence or absence of FTW-mTSCs (unlabeled). Interestingly, after 5 days of co-culture in the FTW condition, many FTW-mESC colonies were surrounded by FTW-mXENs. These FTW-mESC colonies appeared smaller and more “domed” when compared to standalone colonies (not in contact with FTW-mXENs) and FTW-mESC colonies from separate culture (Figure 3B and Video S1). Of note is that we didn’t observe this phenomenon in FTW-mESCs co-cultured with either FTW-mTSCs or mouse embryonic fibroblasts (Figure 3B). In consistent, we found a significant reduction of the GFP signal “Area x Intensity/Colony” in FTW-mESCs co-cultured with FTW-mXENs and FTW-mXENs/FTW-mTSCs (Figure 3C) but not with FTW-mTSCs and fibroblasts, when compared with separately cultured FTW-mESCs. We also calculated the cell density (cell number per cm^2^) of FTW-mESCs daily in each experimental group. On day 5, a significantly lower density of FTW-mESCs was found in FTW-ESCs/TSCs/XENs and FTW-ESCs/XENs groups than in control groups (Figure 3D and Figure S3A). We tested the effects of different cell plating ratios and found greater reduction in FTW-mESC densities when co-cultured with more FTW-mXENs in both 2D and 3D (Figure S3B to S3D). This decrease in FTW-mESC density in co-culture was not due to increased cell apoptosis (Figure S3E and S3F) or differentiation (Figure S3G) and was dependent on direct contact with FTW-mXENs (Figure S3H, S3I and Video S1).

**Figure 3.**
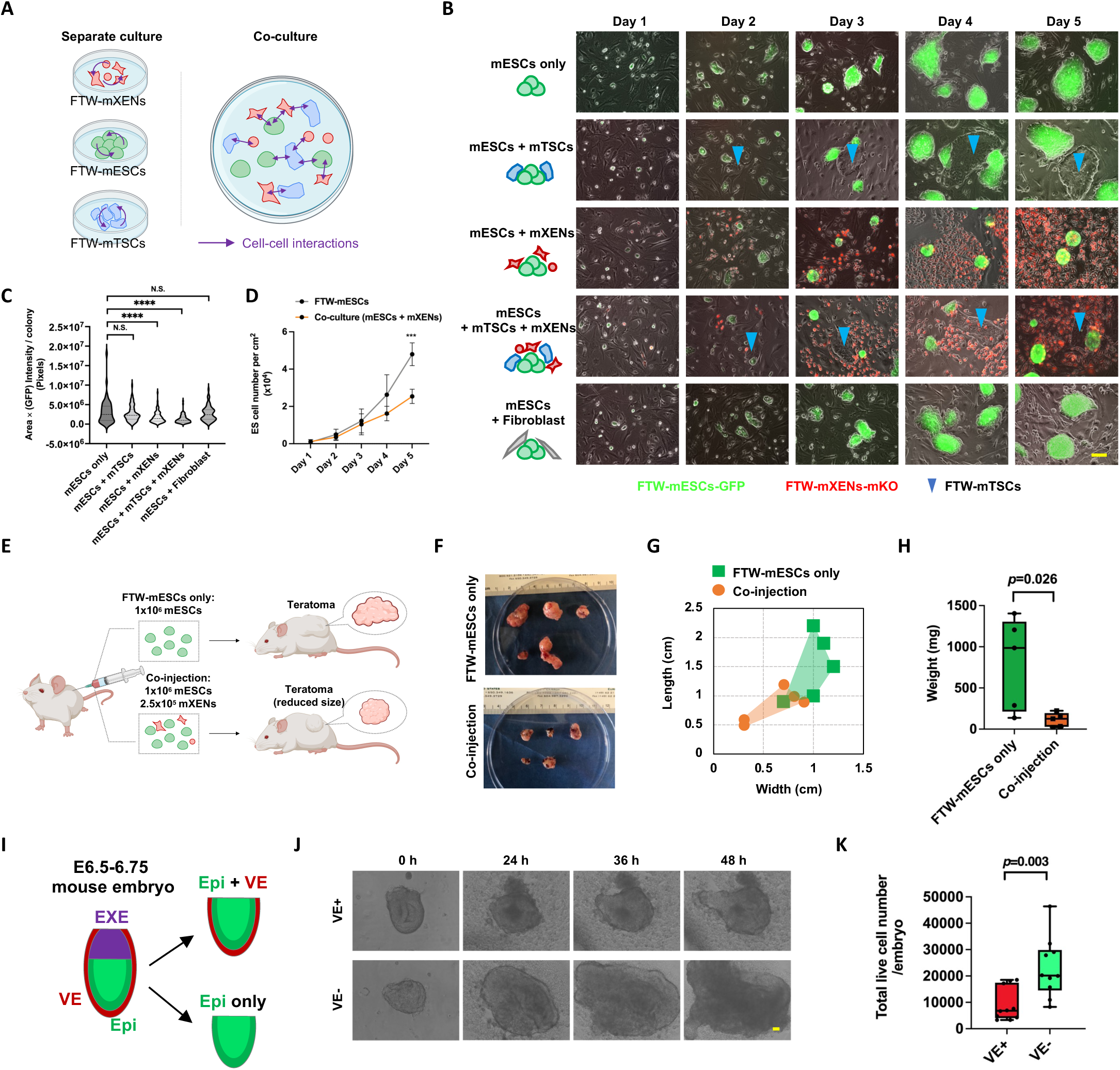
Proliferation restriction of FTW-mESCs by FTW-mXENs. (A) Schematic of the establishment of FTW stem cell co-cultures to study cross-lineage communications. (B) Representative fluorescence and BF merged images of day 1 to day 5 separately cultured FTW-mESCs (green) and FTW-mESCs co-cultured with FTW-mXENs (red) and/or FTW-mTSCs (blue arrowheads), or proliferative mouse embryonic fibroblast. Scale bar, 100 μm. (C) Violin plot showing area multiplied by GFP intensity for each FTW-mESC colonies in day 5 separate and different co-cultures. (D) Growth curves of separate (FTW-mESCs) and co-cultures (mESCs/mXENs) from days 1 to 5 (mean ± SD, day 1, n = 2, day 2-5, n = 5, biological replicates). (E) Schematic of teratoma formation with FTW-mESCs only and FTW-mESCs co-injected with FTW-mXENs (mESCs:mXENs = 4:1). The same number (1 x 10^6^) of FTW-mESCs were injected in each condition. (F) Images of teratomas generated from FTW-mESCs injected with (bottom) and without (top) FTW-mXENs. (G) Lengths and widths of teratomas generated from FTW-mESCs injected with (orange) and without (green) FTW-mXENs. (H) Weight of teratomas generated from FTW-mESCs injected with (orange) and without (green) FTW-mXENs. (mean ± SD, n = 5, biological replicates). (I) Schematic of tissue dissection of E6.5-6.75 mouse conceptus. (J) Representative BF images showing *ex vivo* culture of EPI+VE (VE+) and EPI (VE−) tissues isolated from E6.5-6.75 mouse conceptuses at indicated time points. Scale bar, 100 µm. (K) Total cell number of VE+ and VE− tissues after 48 h *ex vivo* culture (mean ± SD, n = 10, biological replicates). N.S. not significant, ****P-value < 0.0001, P-values were calculated using two-tailed Student’s t test. See also Figure S3.

Next, we studied whether this growth inhibition phenotype manifested during differentiation. We injected the same number of FTW-mESCs either alone or together with FTW-mXENs under the skin of NOD-SCID mice (Figure 3E). Although in both conditions teratomas that contained tissues from all three germ lineages could be generated (Figure S3J), the average size and weight of teratomas generated by co-injection were smaller than FTW-mESCs alone (Figure 3F to 3H). We also tested different ESC:XEN ratios and found in general the more FTW-mXENs were injected, the smaller the teratomas (Figure S3K). To study whether this growth restriction also exists between EPI and VE cells, we isolated E6.5-6.75 mouse conceptuses, removed ExE and ectoplacental cone, and used EPI with or without VE (VE+/−) for *ex vivo* culture (Figure 3I). Interestingly, the size and total cell number in the VE+ group were significantly smaller than those in the VE− group (Figure 3J, 3K and Figure S3L), suggesting the proliferation of EPI is also limited by the VE.

Collectively, we established embryonic and extra-embryonic stem cell co-cultures and identified a contact-dependent growth restriction of FTW-mESCs by FTW-mXENs, which may reflect an embryo size control mechanism during early development.

### Mechanistic insights

To gain mechanistic insights into the crosstalk among co-cultured FTW stem cells, we performed scRNA-seq analysis (Figure 4A). UMAP analysis showed that cells in co-cultures largely overlapped with separately cultured cells (Figure S4A and S4B). We used the CellChat(Jin et al., 2021) toolkit to infer cell-cell communication. As expected, both the number and strength of cell-cell interactions increased in co-cultures when compared to separate cultures (Figure 4B). Interestingly, a significant proportion of these interactions originated from FTW-mXENs towards the other two cell types, especially FTW-mESCs (Figure 4C). CellChat analysis also predicted several signaling pathways mediating these interactions, which included signaling through extracellular matrix (ECM) proteins such as LAMININ and COLLAGEN (Figure 4D). LAMININ and COLLAGEN are known components of the basement membrane (BM) lining the basal side of the post-implantation EPI, which are in part produced by the VE in mice(Sekiguchi and Yamada, 2018). We next determined whether ECM protein(s) could phenocopy the growth restriction of co-cultured FTW-mESCs. We found supplementation of separately cultured FTW-mESCs with Matrigel (~60% LAMININ, ~30% COLLAGEN IV), LAMININ, or COLLAGEN IV, but not VITRONECTIN could inhibit FTW-mESC growth in a dosage dependent manner (Figure 4E, and Figure S4C, S4D). In consistent, knockout of *Laminin* γ*1* (*Lamc1*) in FTW-mXENs could partially rescue the growth phenotype of co-cultured FTW-mESCs (Figure 4F and Figure S4E, S4F). As INTEGRIN-β1 plays an important role in ECM protein signaling and is a cell surface receptor for LAMININ-γ1(Barczyk et al., 2010; Molè, 2021; Moore et al., 2014), we generated *integrin* β*1* (*Itgb1*) knockout FTW-mESCs. We found loss-of-function of *Itgb1*could also mitigate the retarded growth of co-cultured FTW-mESCs (Figure S4G and S4H).

**Figure 4.**
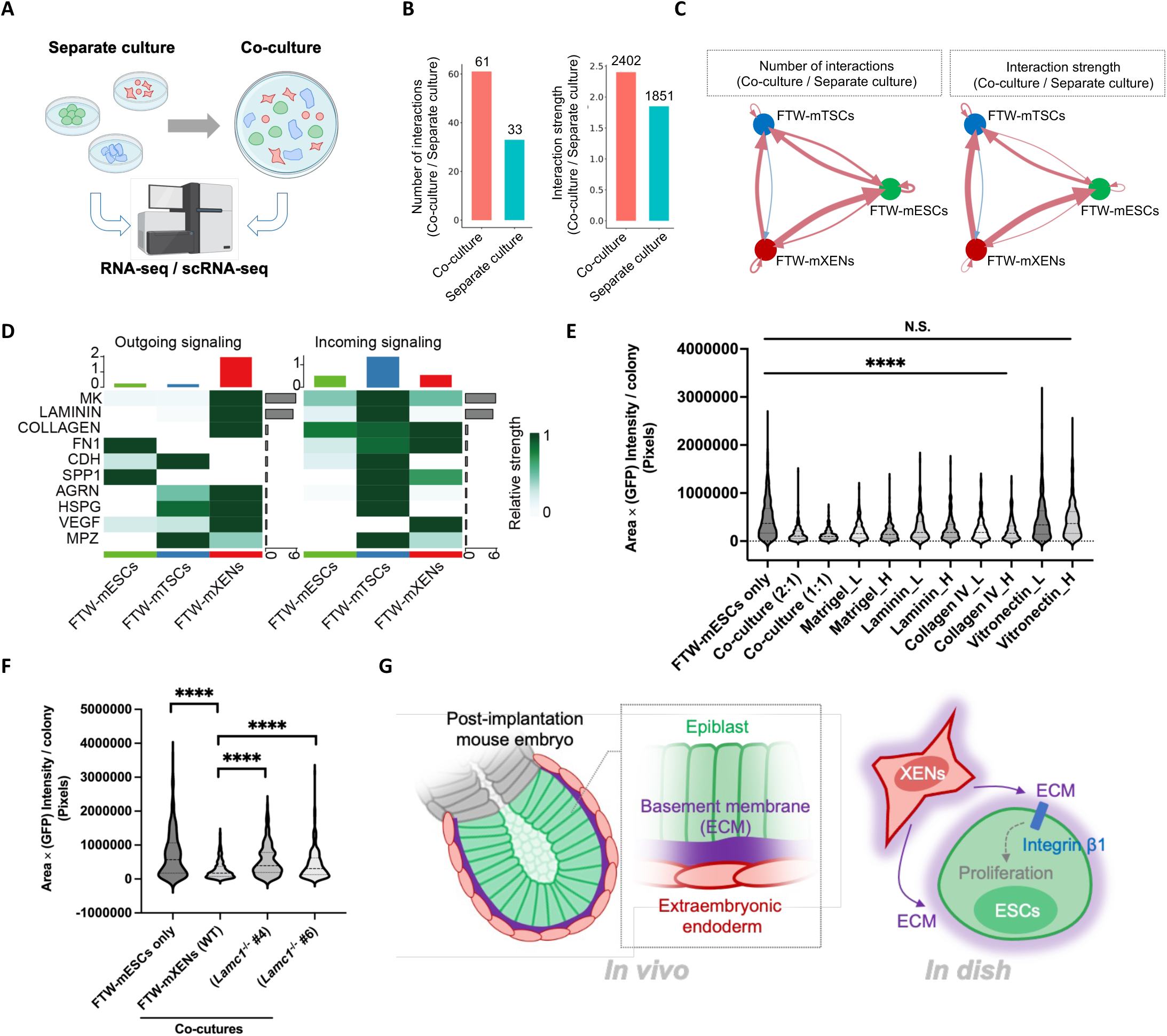
Mechanistic insights of growth inhibition. (A) Schematic of RNA-seq experiments. (B) Bar graphs showing the number (left) and strength (right) of cell-cell interactions in co-cultures and separate cultures. (C) Circle plots showing the ratios of number (left) and strength (right) of cell-cell interactions between co-cultured and separately cultured samples. Red lines, increased interactions; blue lines, decreased interactions. (D) Heatmaps of outgoing signaling patterns (left) and incoming signaling patterns (right) in co-cultured mouse FTW stem cells. (E) Violin plot showing area multiplied by GFP intensity for each FTW-mESC colonies in day 5 separate and co-cultures (mESCs: mXENs = 2:1 or 1:1) as well as separate cultures supplemented with different ECM proteins. Matrigel_L: 0.5% (v/v), Matrigel_H: 2% (v/v), Laminin_L: 30 μg/ml, Laminin_H: 120 μg/ml, Collagen_L: 15 μg/ml, Collagen_H: 60 μg/ml, Vitronectin_L: 5 μg/ml, Vitronectin_H: 30 μg/ml. N.S. not significant. (F) Violin plot showing area multiplied by GFP intensity for each FTW-mESC colonies in different conditions. (G) Schematic summary of mechanistic insights of the observed proliferation inhibition of FTW-mESCs by FTW-mXENs. N.S. not significant, ****P-value < 0.0001. P-values were calculated using two-tailed Student’s t test. See also Figure S4.

In addition, we studied the transcriptomic differences between separately cultured and co-cultured FTW-mESCs by bulk RNA-seq. Through comparative analysis, we identified 502 differentially expressed genes (DEGs) shared between FTW-mESCs in XENs/ESCs and XENs/TSCs/ESCs co-cultures, when compared to separately cultured FTW-mESCs (Figure S4I and Table S2). Interestingly, the majority of DEGs (492) were down-regulated genes in co-cultured FTW-mESCs. Consistent with their decreased proliferation, the down-regulated DEGs included genes related to cell proliferation and embryo size (e.g., several members of activator protein 1[AP-1] transcription factor complex such as *Fos*, *Fosl1*, *Fosb*, *Batf* among others(Angel et al., 1987; Angel and Karin, 1991; Jochum et al., 2001))(Figure S4J). GO and Bioplanet 2019 analyses of the down-regulated DEGs revealed top enriched terms including extracellular matrix and structure organization, Interleukin-1 and TGF-beta regulation of extracellular matrix, and β1 integrin cell surface interactions (Figure S4K and S4L). In addition, we found many matrix metalloproteinases (MMPs) (e.g., *Mmps 2, 3, 9, 10, 12* etc.) were down-regulated in co-cultured FTW-mESCs (Figure S4M). MMPs have been reported to modulate cell proliferation, migration, and morphogenesis by degrading ECM proteins(Vu, 2000), and inhibition of MMPs also led to reduced growth of EPI in vivo and ESCs in vitro(Kyprianou et al., 2020), which are consistent with our findings.

In sum, these analyses helped gain insights into the crosstalk among co-cultured embryonic and extraembryonic stem cells and identified ECM signaling as one of the mechanisms mediating the reduced proliferation of FTW-mESCs by FTW-mXENs (Figure 4G).

### Stem cells from monkey blastocysts

By using the FTW condition, we also succeeded in the derivation of several stable extra-embryonic (FTW-cyXENs and FTW-cyTSCs) and embryonic (FTW-cyESCs, 20% KSR [knockout serum replacement] was needed) stem cell lines from 7-10 d.p.f. (days post-fertilization) cynomolgus monkey blastocysts (Figure 5A to 5D, and Table S3). FTW-cyESCs could also be directly converted from naïve-like ESCs(Fang et al., 2014) through culture adaptation (data not shown). Once established, FTW-cyXENs, FTW-cyTSCs and FTW-cyESCs proliferated well, maintained stable colony morphology and normal karyotypes after long-term culture, expressed hypoblast (HYP), trophoblast, and EPI related genes, respectively (Figure 5E and Figure S5A to S5D, S5E to S5H, S5L, S5M). Upon random differentiation *in vitro*, FTW-cyXENs could generate visceral- and yolk-sac-endoderm- (VE/YE-) (FOXA1^+^ GATA4^−^) like cells and extra-embryonic-mesenchyme-cell- (EXMC−) (COL6A1^+^ GATA4^−^) like cells(Nakamura et al., 2016; Niu et al., 2019b) (Figure 5F and S5I). FTW-cyTSCs were capable of differentiating into extravillous trophoblast (EVT)-like cells and multinucleated syncytiotrophoblast (SCT)-like cells *in vitro* (Figure 5G and 5H). (Okae *et al*., 2018) and in TSC-teratomas (Figure S5J and S5K).

**Figure 5.**
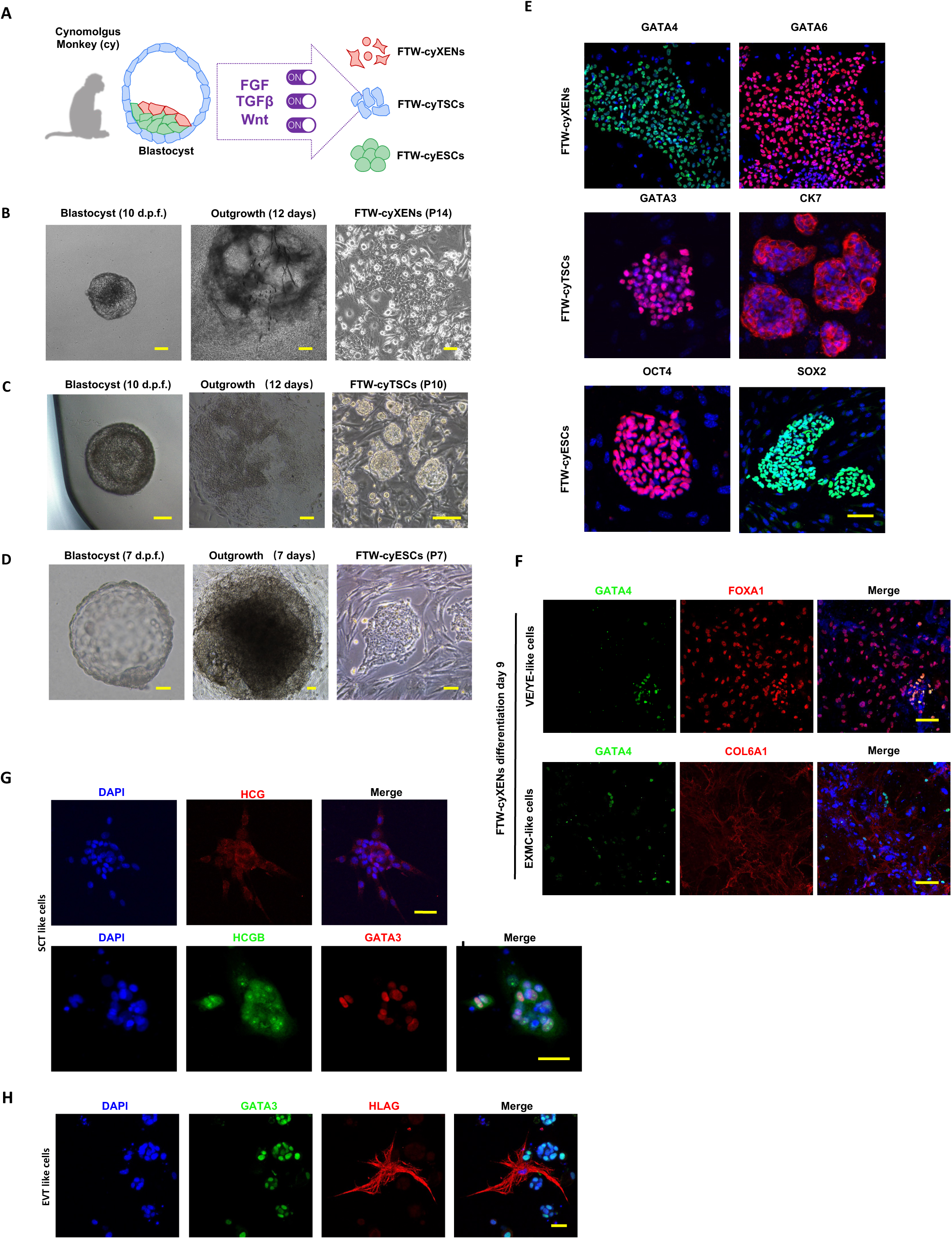
Cynomolgus monkey embryonic and extraembryonic stem cells derivation. (A) Schematic of derivation of FTW embryonic and extra-embryonic stem cell lines from cynomolgus monkey blastocyst. (B) Representative BF images of a 10 d.p.f monkey blastocyst, day 12 outgrowth and stable FTW-cyXENs (passage 14). Scale bars, 50 µm. (C) Representative BF images of a 10 d.p.f monkey blastocyst, day 12 outgrowth and stable FTW-cyTSCs (passage 10). Scale bars, 50 µm. (D) Representative BF images of a 7 d.p.f monkey blastocyst, day 7 outgrowth, FTW-cyESCs, passage 7. Scale bars, 50 µm. (E) Representative IF images of monkey extra-embryonic endoderm (GATA6 and GATA4), trophoblast (GATA3 and CK7), and epiblast (OCT4 and SOX2) lineage markers in FTW-cyXENs (top), FTW-cyTSCs (middle), and FTW-cyESCs (bottom), respectively. Scale bar, 100 µm. (F) Representative IF co-staining images of COL6A1, FOXA1, and GATA4 in differentiated FTW-cyXENs at day 9. Blue, DAPI. Scale bars, 100 µm. (G) Representative IF co-staining images of GATA3 with the EVT maker HLAG in SCT-like cells differentiated from FTW-cyTSCs. (H) Representative IF co-staining images of GATA3 with the EVT makers HCG and HCGB in EVT-like cells differentiated from FTW-cyTSCs. See also Figure S5.

Next, we performed transcriptomic profiling of cynomolgus monkey FTW stem cells. scRNA-seq analysis revealed that FTW-cyESCs, FTW-cyXENs and FTW-cyTSCs segregated into three distinct clusters on UMAP and expressed their respective lineage marker genes (Figure 6A and 6B). Bulk RNA-seq analysis helped identify DEGs among FTW-cyESCs, FTW-cyXENs and FTW-cyTSCs, which were enriched in GO terms including: “Signaling pathways regulating pluripotency of stem cells” and “VEGF signaling pathway” (FTW-cyESCs); “ECM-receptor interaction” and “TGF-beta signaling” (FTW-cyXENs); “ECM-receptor interaction” and “Hippo signaling pathway” (FTW-cyTSCs) (Figure 6C). The result of the Pearson correlation analysis showed that FTW-cyESCs, FTW-cyXENs and FTW-cyTSCs shared the highest correlation coefficients with their *in vivo* counterparts (Figure 6D).

**Figure 6.**
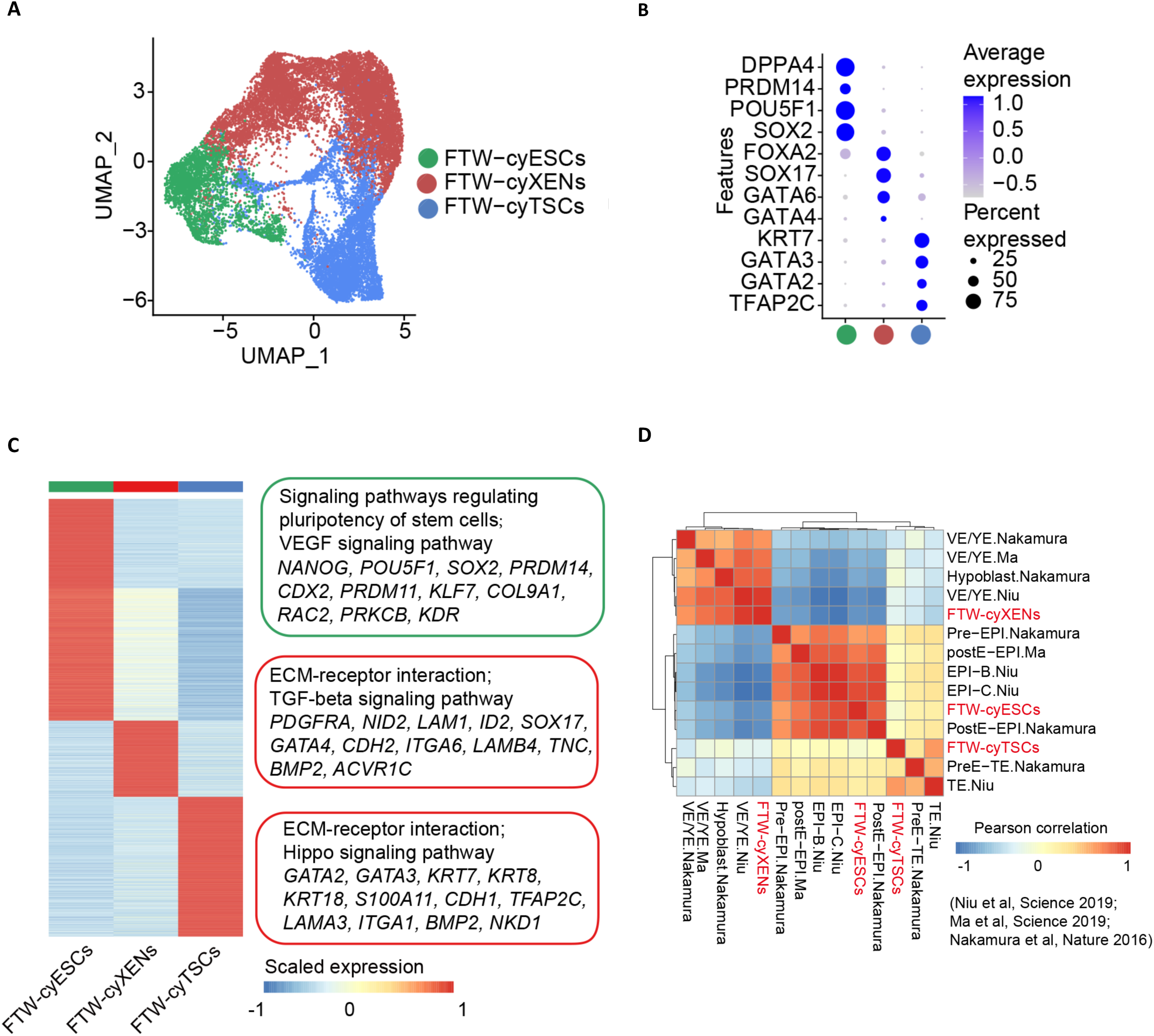
Transcriptomic analyses of cynomolgus monkey FTW stem cells. (A) UMAP analysis of FTW-cyXENs, FTW-cyTSCs and FTW-cyESCs. (B) Bubble plot showing the expression patterns of lineage makers in FTW-cyXENs, FTW-cyTSCs and FTW-cyESCs. (C) A heatmap showing the DEGs of FTW-cyESCs, FTW-cyXENs and FTW-cyTSCs, Selected genes from the DEGs and top two enriched GO terms were shown on the right. (D) Correlation analysis of FTW-cyXENs, FTW-cyTSCs, FTW-cyESCs and published datasets from cynomolgus monkey conceptuses.

### Cross-lineage stem cell co-cultures among monkey cells

Next, we investigated whether the growth restriction of FTW-ESCs by FTW-XENs was conserved in cynomolgus monkeys. To this end, we performed co-culture of FTW-cyESCs with FTW-cyXENs and/or FTW-cyTSCs. Consistent with mouse findings, we found the growth of FTW-cyESCs was greatly inhibited by FTW-cyXENs but marginally affected by FTW-cyTSCs or fibroblasts (Figure 7A, 7B and Figure S6A) To provide mechanistic insights into embryonic and extra-embryonic lineage crosstalk in cynomolgus monkeys, we performed scRNA-seq analysis of co-cultures of FTW-cyESCs/FTW-cyXENs (2lines) and FTW-cyESCs/FTW-cyXENs/FTW-cyTSCs (3lines) using 10x Genomics Chromium, and compared with single cell transcriptomes derived from separate cultures (Table S4). Through DEG and GO analyses, we identified several terms, e.g., “Focal adhesion”, “Cadherin binding”, and “Cell-cell adhesion”, were enriched in co-cultured FTW-cyESCs, FTW-cyXENs and FTW-cyTSCs, respectively, suggesting more active cell-cell interactions (Figure S6B to S6D). In agreement with mouse results, CellChat analysis also predicted signaling through ECM proteins, e.g., LAMININ and COLLAGEN, mediated crosstalk between FTW-cyXENs and FTW-cyESCs (Figure 7C, Figure S6E and Table S5). To confirm the effect(s) of ECM protein(s), we supplemented Matrigel, LAMININ, or COLLAGEN IV to separately cultured FTW-cyESCs, and found that each could phenocopy FTW-cyXENs’ inhibitory effect on the growth of FTW-cyESCs in a dosage dependent manner (Figure 7D and Figure S6F, S6G).

**Figure 7.**
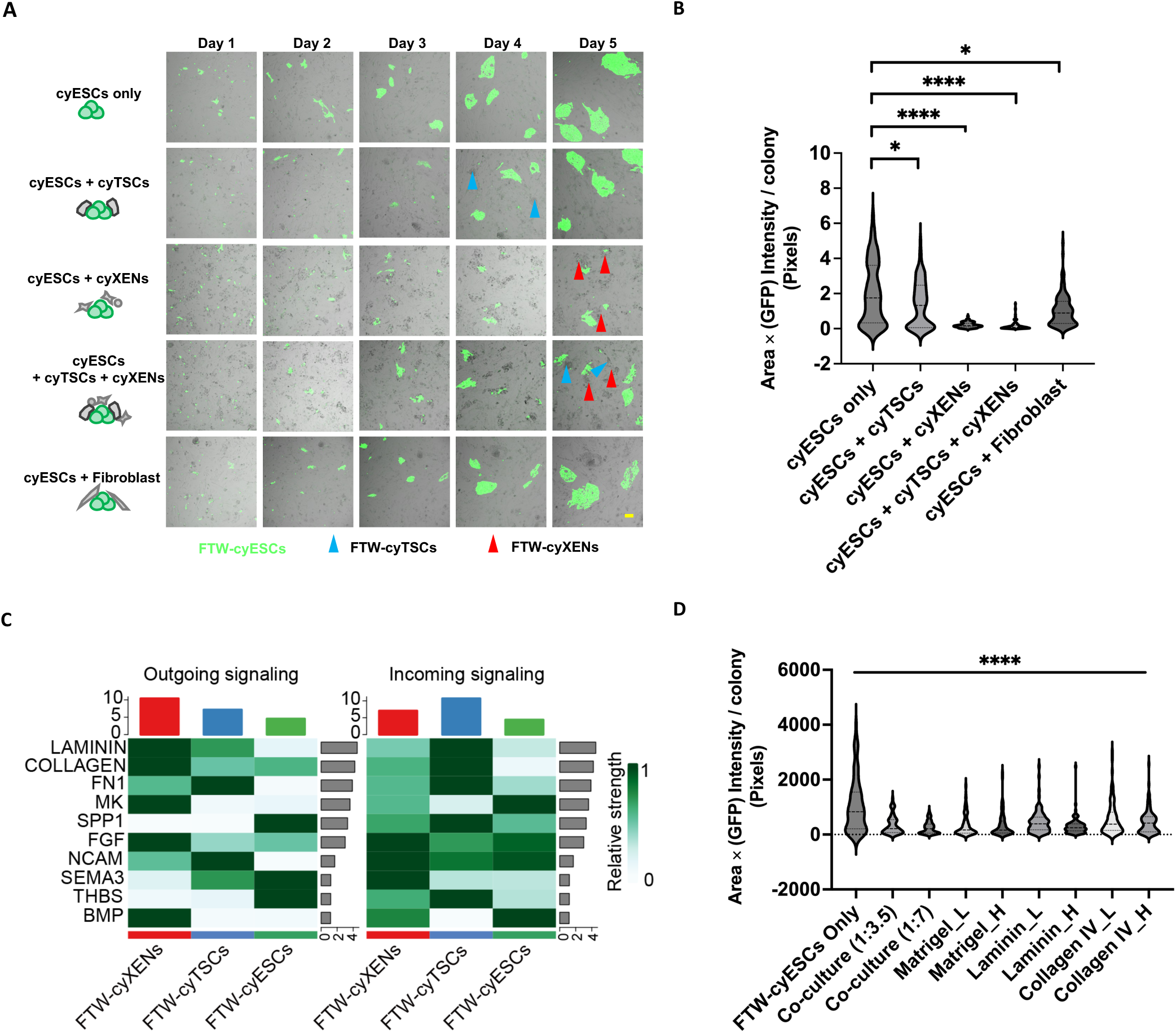
Cynomolgus monkey embryonic and extraembryonic stem cells co-cultures. **(A)** Representative fluorescence and BF merged images of day 1 to day 5 separately cultured FTW-cyESCs (green) and FTW-mESCs co-cultured with FTW-cyXENs (red arrowheads) and/or FTW-cyTSCs (blue arrowheads), or proliferative fibroblast. Scale bar, 100 μm. **(B)** Violin plot showing area multiplied by GFP intensity for each FTW-mESC colonies in different conditions. **(C)** Heatmaps of outgoing signaling patterns (left) and incoming signaling patterns (right) in co-cultured cynomolgus FTW stem cells. **(D)** Violin plot showing area multiplied by GFP intensity for each FTW-cyESC colonies in day 5 separate and co-cultures (cyESCs: cyXENs = 1:3.5 or 1:7) as well as separate cultures supplemented with different ECM proteins. Matrigel_L: 0.5% (v/v), Matrigel_H: 2% (v/v), Laminin_L: 2.5 μg/ml, Laminin_H: 10 μg/ml, Collagen_L: 15 μg/ml, Collagen_H: 60 μg/ml. *P-value < 0.05, ****P-value < 0.0001. P-values were calculated using two-tailed Student’s t test. See also Figure S6.

Taken together, these results extended our findings in mice to cynomolgus monkeys and revealed a conserved mechanism underlying growth inhibition of FTW-ESCs by FTW-XENs.

### Cross species comparisons

Embryonic and extraembryonic stem cells from both mice and cynomolgus monkeys derived and cultured in the same condition gave us a unique opportunity to examine species-conserved and -divergent features not influenced by different culture parameters. Using the single-cell regulatory network inference and clustering (SCENIC) pipeline, we first studied the regulatory networks underlying the maintenance of FTW-ESCs, FTW-XENs and FTW-TSCs in both species. Our analysis identified several conserved and species-specific transcription factor (TF)-driven regulons in each stem cell type (Figure S7A to S7C). SCENIC analysis also revealed common and distinct targets of well-known lineage TFs such as NANOG, SOX17 and GATA3 (Figure S7A to S7C and Table S6). We next performed cross-species comparison of intercommunications between lineages, which uncovered both species-conserved (e.g., MK, LAMININ, COLLAGEN and FN1) and -divergent (e.g., CDH, AGRN and HSPG [mouse]; FGF, NCAM and BMP [monkey]) signaling crosstalk (Figure S7D). Collectively, these results reveal interspecies transcriptomic similarities and differences in early embryonic and extraembryonic cells and their intercommunications, which provide insights into evolutionary convergent and divergent processes underlying epiblast, hypoblast and trophoblast development across different species *in vivo*.

## DISCUSSION

Most stem cell cultures include pathway inhibitors to suppress differentiation in favor of self-renewal, e.g., ground state mouse ESC culture (Ying et al., 2008), which often necessitate the design of new conditions for different stem cell types and different species. By activating multiple developmental signaling pathways at once, surprisingly, we found several embryonic (mouse, cynomolgus monkey, and horse) and extraembryonic stem cells (mouse and cynomolgus monkey) from multiple species could be derived and stably maintained in the same condition (Wu et al., 2017; Yu *et al*., 2021b). This signaling “excited” state can potentially be attained in a variety of stem cell types by striking a balance between differentiation and self-renewal via switching on a combination of signaling pathways including but not limited to FGF, TGF-β and WNT pathways used in this study.

The ability to grow embryonic and extraembryonic stem cells in the same culture environment opens new avenues to dissect the molecular and cellular mechanisms underlying lineage crosstalk during early mammalian development. Based on FTW stem cell co-cultures, we have uncovered a previously unrecognized non-cell autonomous control of pluripotent cell proliferation by extraembryonic endoderm cells, which is in part mediated by the ECM signaling, in both mice and cynomolgus monkeys. We further revealed a similar overgrowth phenotype in mouse epiblast in the absence of visceral endoderm, providing evidence supporting the in vivo relevance of the observed proliferation restriction. Consistant with this finding, it was previously reported that MMP-mediated basement membrane perforations at the posterior side of the mouse embryo permited growth and primitive-streak extension during peri-gastrulation development (Kyprianou *et al*., 2020).

Our study identified laminin-integrin signaling contributed to the growth restriction of FTW-mESCs by FTW-mXENs (Figure 4D-4F and Figure S4C-S4F). This is likely due to irregularly distributed matrix materials surrounding integrin β1-deficient FTW-mESCs during co-culture with FTW-mXENs as opposed to a continuous basement membrane layer in WT FTW-mESCs (Moore *et al*., 2014). A previous study demonstrated that integrin β1-deficient mESCs underwent massive cell death during transition from naive pluripotency toward the formative pluripotency (Molè, 2021). This seemingly contrasts with integrin β1-deficient formative FTW-mESCs (Figures S4G, S4H). A possible explaination is that despite FTW-mESCs exhibited functional formative features including the ability to form blastocyst chimeras and direct responsiveness to primordial germ cell specification, they retain the expression of many naïve pluripotency genes and are recognized as more naïve-like formative PSCs as apposed to more primed-like formative stem (FS) cells (Kinoshita et al., 2021; Yu *et al*., 2021b). In consistent, the induction of of apoptosis of integrin β1-deficient mESCs during differetnation was found concomitant with the upregulation of OTX2, and OTX2 level was found low in FTW-mESCs (Molè, 2021).

The recent advancements in both extended primate (human and cynomolgus monkey) embryo cultures and stem cell embryo models provide an unprescendented opportunity to study primate early post-implantation development (Deglincerti et al., 2016; Ma et al., 2019; Moris et al., 2020; Niu et al., 2019a; Shahbazi et al., 2016; Yu et al., 2022; Yu et al., 2021a; Zheng et al., 2019). However, the dynamic nature of a developing embryo/embryo model and its cellular complexity make it difficult to dissect the key cell-cell interactions. Here, the establishment of cynomolgus monkey FTW-ESCs, XENs and TSCs under the same condition enables us, for the first time, to delineate cell-cell interactions in a tractable and accessible manner in primates. In future studies, the combination of stem cell co-culture with CRISPR screens will help uncover additional genes and pathways underlying the crosstalk between extraembryonic and embryonic cells during embryogenesis in primates.

In sum, the FTW embryonic and extraembryonic stem cells and the co-culture strategy developed in this study may help provide superior starting cells for generating more robust integrated stem cell embryo models (Fu et al., 2021) and developing more faithful differentiation protocols for regenerative medicine.

## ACKNOWLEDGMENTS

We thank Janet Rossant for providing the GFP labelled conventional mouse XEN and TSCs. We thank Ling Zhang for technical support. We thank all other members of the Wu laboratory for discussion and suggestions. We also acknowledge UT Southwestern Genomics and Microarray Core for providing next-generation sequencing services. J.W. is a New York Stem Cell Foundation–Robertson Investigator and Virginia Murchison Linthicum Scholar in Medical Research and supported by CPRIT (RR170076), NIH (GM138565-01A1, HD103627-01A1, R21HD105349 and OD028763), ARSM Research Institute, and Welch (854671). T. T. is supported by grants from the National Natural Science Foundation of China (82192871), the National Key Research and Development Program (2021YFA0805700) and the Major Basic Research Project of Science and Technology of Yunnan (202001BC070001 and 202102AA100053). L. X. is founded by the Rally Foundation, The National Institutes of Health (R21CA259771, UM1-HG011996, R01CA263079, R01DK127037, and R01HL144969), CPRIT (RP220032, RP180319, RP200103, and RP180805).

## AUTHOR CONTRIBUTIONS

J.W. T.T. Y.W. and L.Y. conceptualized the idea, designed, analyzed, interpreted the experimental results. Y.W. and L.Y. performed most of the mouse cell culture experiments. E.Z. performed most of the monkey cell culture experiments. B.C. L.G. J.L. H.S. L. X. and T.T. performed RNA-seq analysis. S.T. and D.O. performed mouse E6.5-6.75 embryo with/remove EXE in vitro culture experiment. Y.D. helped with data analysis. C.Z. helped with some mouse cell culture experiments. L.Z. helped with some monkey cell culture experiments. D. S. designed and prepared some DNA constructs. Y.W. and M.S. performed chimera experiments. J.W. T.T. and W.J. supervised the study. Y.W. L.Y. T.T. and J.W. wrote the manuscript with inputs from all authors.

## DECLARATION OF INTERESTS

Y.W., L.Y., T.T. and J.W. are inventors on a patent application (applied through the Board of Regents of The University of Texas System, application number 63/488,401) entitled “Methods For the Derivation Culture Of Embryonic and Extra-Embryonic Stem Cells” arising from this work. The other authors declare no competing interests.

## Data availability

A total of thirty-one RNA-seq datasets generated in this study have been deposited in the Gene Expression Omnibus (GEO) with accession code GSE178459.

## Code availability

The sequencing data generated in this study have been deposited to the Gene Expression Omnibus (GEO) under the accession (GSE178459). They will be made publicly accessible upon publication.

## SUPPLEMENTAL INFORMATION

**Figure S1. Derivation, characterization of FTW-mXENs, FTW-mTSCs and FTW-mESCs and developmental potential of FTW-mXENs and FTW-mTSCs.**

**Figure S2. Transcriptomic analyses of mouse FTW stem cells.**

**Figure S3. Proliferation restriction of FTW-mESCs by FTW-mXENs.**

**Figure S4. Mechanistic insights of growth inhibition and transcriptomic analyses of co-cultured FTW stem cells.**

**Figure S5. Derivation and characterization of FTW-cyXENs, FTW-cyTSCs and FTW-cyESCs.**

**Figure S6. Lineage crosstalk among co-cultured cynomolgus monkey FTW stem cells.**

**Figure S7. Cross-species comparison.**

## METHOD DETAILS

### Mice

C57BL/6 and CD-1 (ICR) mice were purchased from Charles River or Envigo (Harlen). NOD-SCID (NOD.CB17-*Prkdc^scid^*/J) mice were purchased from the Jackson Lab. Mice were housed in 12-hr light/12-hr dark cycle at 22.1–22.3°C and 33–44% humidity. All procedures related to animals were performed in accordance with the ethical guidelines of the University of Texas Southwestern Medical Center. The animal protocol was reviewed and approved by the UT Southwestern Institutional Animal Care and Use Committee (IACUC) [Protocol #2018-102430]. All experiments followed the 2021 Guidelines for Stem Cell Research and Clinical Translation released by the International Society for Stem Cell Research (ISSCR). Human-mouse chimeric studies were reviewed and approved by the UT Southwestern Stem Cell Oversight Committee (SCRO) [Registration #14].

### Cynomolgus Monkeys

All animals and experimental procedures were approved in 2021 by the ethical committee of the LPBR and Institute of Primate Translational Medicine and Kunming University of Science and Technology (IPTM, KUST), and performed by following the guidelines of the Association for Assessment and Accreditation of Laboratory Animal Care International (AAALAC) for the ethical treatment of non-human primates. 2 healthy female cynomolgus monkeys (Macaca fascicularis), ranging in age from 5 to 8 years with body weights of 4 to 6 kg, were selected for use in this study. All animals were housed at the State Key Laboratory of Primate Biomedical Research (LPBR). All cynomolgus monkey embryos related work was conducted at the State Key Laboratory of Primate Biomedical Research.

### Harvesting and culture of mouse embryos

C57BL/6 female mice (8–10 weeks old) were super ovulated by intraperitoneal (IP) injection with 5 IU of PMSG (Prospec), followed by IP injection with 5 IU of hCG (Sigma-Aldrich) 48 h later. After mating with C57BL/6 male mice, embryos at 8-cell to morula stages were harvested at E2.75 [the presence of a virginal plug was defined as embryonic day 0.5 (E0.5)) in KSOM-HEPES by flushing oviducts and uterine horns. Embryos were cultured in the mKSOMaa overnight until blastocyst stage in a humidified atmosphere containing 5% (v/v) CO_2_ and 5% (v/v) O_2_ at 37 °C. CD-1 females (8 weeks old or older) in natural estrous cycles were mated with CD-1 males. Blastocysts were harvested at E3.5 by flushing uterine horns.

### Derivation and culture of FTW-mXENs, FTW-mTSCs and FTW-mESCs

Embryo manipulations were performed under a dissection microscope (Nikon SMZ800N). In brief, zona pellucidae (ZP) were removed from E3.5 blastocysts by brief treatment with acidic Tyrode’s solution (Millipore MR-004-D). After removing zona pellucidae, embryos were plated on MEFs in FTW medium [N2B27 basal medium supplemented with FGF2 (20 ng/mL, Peprotech), Activin-A (20 ng/mL, Peprotech) and CHIR99021 (3 μM, Selleckchem). After 6-8 days in culture, blastocyst outgrowths were passaged using TrypLE and re-seeded onto newly prepared MEFs. XEN, TSC, and ESC colonies were manually picked for further cultivation. Established mouse XEN, TSC, and ESC lines were cultured on MEFs plates pre-coated with 0.1% gelatin in FTW medium. The cells were cultured at 37 °C under 5% CO_2_, with daily media changes. For passaging, the cells were dissociated into single cells using TrypLE Express (GIBCO) and passaged onto new MEF-coated plates at a split ratio of 1:20 (FTW-mESCs), 1:10 (FTW-mTSCs), and 1:50 (FTW-mXEN) every 3–4 days.

### Generation of fluorescent mouse FTW cells

We used pCAG-IP-mKO or pCAG-IP-eGFP to label FTW-mXENs,FTW-mTSCs, and FTW-mESCs. In brief, 1–2 μg of pCAG-IP-mKO/eGFP plasmids were transfected into 1 × 10^6^ – 2 × 10^6^ dissociated single cells using an electroporator (NEPA21, Nepa Gene) following the protocol recommended by the manufacturer. Then, 0.5–1.0 μg ml^−1^ of puromycin (Invitrogen) was added to the culture medium 2–3 days after transfection. Drug-resistant colonies were manually picked between 7 to 14 days and further expanded clonally.

### *In vitro* differentiation of FTW-mTSCs and FTW-mXENs

FTW**-**XENs were dissociated into single cells using TrypLE Express and seeded into a 6-well plate pre-coated with 5 μg/mL Laminin at a density of 2 × 10^5^ cells (mouse XEN) or 5 × 10^5^ cells (monkey and human XEN) per well in XEN differentiation medium for 9 days, with medium changes every other day. XEN differentiation medium was prepared using the following: 1:1 (v/v) mixture of DMEM/F12 and Neurobasal medium, 1X N2 supplement, 1X B27 minus insulin supplement, 1X GlutaMAX, 1X Nonessential amino acids, 0.1 mM β-mercaptoethanol, 0.5% Penicillin-Streptomycin, and 10% FBS.

### Harvesting and culture of cynomolgus embryos

Cynomolgus monkey (Macaca fasicularis) ovarian stimulation, oocyte recovery and in vitro fertilization were performed as previously described(Niu et al., 2014). Briefly, healthy female cynomolgus monkeys were subjected to follicular stimulation by intramuscular injection of 20 IU of recombinant human follitropin alpha (rhFSH, Gonal F, Merck Serono) for 8 days, then 1,000 IU recombinant human chorionic gonadotropin alpha (rhCG, OVIDREL, Merck Serono) was injected on day 9. Cumulus-oocyte complexes were collected by laparoscopic follicular aspiration 32-35 hours following rhCG administration. Follicular contents were placed in HEPES-buffered Tyrode’s albumin lactate pyruvate (TALP) medium containing 0.3% bovine serum albumin (BSA) (Sigma-Aldrich) at 37°C. Oocytes were stripped of cumulus cells by pipetting after a brief exposure (<1 min) to hyaluronidase (0.5 mg/mL) in TALP-HEPES to allow visual selection of nuclear maturity metaphase II (MII; first polar body present) oocytes. The maturity oocytes were subjected to intracytoplasmic sperm injection (ICSI) immediately and then cultured in CMRL-1066 medium (Gibco, 11530037) containing 10% FBS at 37°C in 5% CO2. Fertilization was confirmed by the presence of the second polar body and two pronuclei. Zygotes were then cultured in the chemically defined hamster embryo culture medium-9 (HECM-9) containing 10% fetal bovine serum at 37 °C in 5% CO2 to allow embryo development. The blastocysts were collected at 7 days post fertilization (d.p.f.). The zona pellucida of blastocyst was removed by exposure to hyaluronidase from bovine testes (Sigma-Aldrich) about 30 seconds and embryos were cultured in vitro until 10 days post fertilization.

### Derivation and culture of FTW-cyXENs, FTW-cyTSCs, FTW-cyESCs

For the derivation of FTW-cyXENs and FTW-cyTSCs, 10 d.p.f. cynomolgus monkey embryos were dissected with a 31-gauge syringe needle and plated into wells of 4-well dish coated with 0.1% gelatin and covered with mitotically inactivated MEF in FTW medium (done the same way as with mouse). To promote the proliferation of hypoblast, 1% KSR (Thermo Fisher Scientific, A3181502) and PDGF (10 ng/mL, R&D, 220-BB) were also added(Zhong *et al*., 2018). After 8-10 days, XEN-like and TS-like colonies appeared. Single colonies were isolated and then dissociated by TrypLE (Thermo Fisher Scientific,12604021) for 3 minutes at 37 °C, and passaged into a well of a 4-well dish. Established FTW-cyXENs and FTW-cyTSCs were passaged by TrypLE every 6-7 days (monkey) at a split ratio of 1:10 in FTW medium under 20% O_2_ and 5% CO_2_ at 37. To derive FTW-cyESCs(Kang et al., 2018), blastocysts (7 days p.c.) were plated into a well of 4-well dish coated with 0.1% gelatin and MEFs in FTW cyESC medium [N2B27 basal medium supplemented with 20% KSR, FGF2 (6 ng/mL, Peprotech), Activin-A (25 ng/mL, Peprotech) and CHIR99021 (1.5 μM, Selleckchem)] under 5% O_2_ and 5% CO_2_. After 8-10 days, the ES-like outgrowths appeared and were dissociated using TrypLE for 3 minutes at 37 °C. The established FTW-cyESCs were cultured in a 6-well plate pre-coated with 0.1% gelatin MEFs in FTW cyESC medium under 20% O_2_ and 5% CO_2_ at 37.

### *In vitro* Differentiation of FTW-cyXENs

To induce differentiation, FTW-cyXENs were dissociated into single cells using TrypLE and seeded into a 6-well plate pre-coated with 5 μg/mL Laminin at a density of 5 × 10^5^ cells per well and kept in differentiation medium for 9 days with medium changes every other day. Differentiation medium was prepared as following: 1:1 (v/v) mixture of DMEM/F12 and Neurobasal medium, 0.5X N2 supplement, 0.5X B27 supplement, 1% GlutaMAX, 1% Nonessential amino acids, 0.1 mM β-Mercaptoethanol, 1% Penicillin-Streptomycin, and 10% FBS.

### *In vitro* Differentiation of FTW-cyTSCs

FTW-cyTSCs were differentiated into ST and EVT as the described with human cells(Okae *et al*., 2018). For the induction of EVT cells, FTW-cyTSCs were dissociated into single cells in a 6-well plate pre-coated with 1 mg/ml Collagen-IV at a density of 1 × 10^5^ cells per well and cultured with 3 mL of EVT medium (DMEM/F12 supplemented with 1% Penicillin-Streptomycin, 0.3% BSA, 4% KnockOut Serum Replacement, 0.1 mM β-Mercaptoethanol, 1% ITS-X supplement, 100 ng/ml NRG1, 2.5 mM Y27632, 7.5 mM A83-01). 2% Matrigel was added to the medium on the first day. At day 3, the medium was changed to EVT medium with 0.5% Matrigel, but without NRG1. At day 6, the medium was replaced with EVT medium with 0.5% Matrigel, but without NRG1 and KSR, and cells were cultured for an additional two days. For differentiation of ST (2D) cells, FTW-cyTSCs were dissociated in a 6-well plate pre-coated with 2.5 mg/ml Collagen-IV at a density of 2 × 10^5^ cells per well and cultured in 3 mL of ST(2D) medium (DMEM/F12 supplemented with 0.1 mM β-Mercaptoethanol, 1% Penicillin-Streptomycin, 4% KSR, 0.3% BSA, 1% ITS-X supplement, 2.5 mM Y27632 and 2 mM Forskolin). The medium was replaced on day 3, and the cells were analyzed on day 6. For the differentiation of ST (3D) cells, 2 × 10^5^ FTW-cyTSCs were seeded in a 3.5 cm Petri dish and cultured with 3 mL of ST (3D) medium (DMEM/F12 supplemented with 0.1 mM β-Mercaptoethanol, 1% Penicillin-Streptomycin, 4% KSR, 0.3% BSA, 1% ITS-X supplement, 2 mM Forskolin, 50 ng/ml EGF, and 2.5 mM Y27632]. An equal amount of fresh ST (3D) medium was added at day 3. The cells were passed through a 40 μm mesh filter to remove dead cells and debris at day 6.

### RT-PCR and qRT-PCR Analysis

Total RNA was isolated using the RNeasy Mini Kit (QIAGEN) following the manufacturer’s instructions. Contaminating genomic DNA was removed by RNase-Free DNase Set (QIAGEN). RNA concentrations were measured on a spectrophotometer (DS-11+, DeNovix). cDNA was synthesized with iScript Reverse Transcription Supermix kit (BIO-RAD) and amplified with PrimeSTAR GXL DNA Polymerase (TaKaRa) or with SYBR Green PCR Master Mix (Thermo Fisher Scientific) on a Touch Thermal Cycler Real-Time PCR system (C1000, BIO-RAD). GAPDH was used as an internal normalization control. All primers used in this study were listed in Table S7.

### Cell population doubling time

The cell population doubling time was calculated using the doubling time online calculator (http://www.doubling-time.com/compute.php?lang=en).

### Cell Growth Curve

FTW-cyXENs, FTW-cyTSCs and FTW-cyESCs’s (1 × 10^5^) were seeded onto 6-well plates coated with MEF in FTW medium, with media changes every day. Cells were harvested by TrypLE Express at the indicated time points, depleted of MEFs by plating the cell suspension onto 0.5% gelatin-coated plates for 30 minutes, and growth curves were generated by manual cell counting.

### Teratoma formation

For FTW-mTSC and FTW-mXEN teratomas, a total of 5 × 10^6^ cells were resuspended in 100 μL of DMEM-Matrigel solution (1:1) and injected subcutaneously into 10-week-old immunodeficient NOD-SCID mice. After 2 weeks, FTW-mTSC teratomas were dissected and fixed with PBS containing 4% formaldehyde. After 3 months, FTW-mXEN teratomas were dissected and fixed with PBS containing 4% formaldehyde. For cyTSC teratomas, a total of 1 × 10^7^ cells were resuspended in 100 μL of DMEM-Matrigel solution (1:1) and injected subcutaneously into 10-week-old immunodeficient NOD-SCID mice. After 7 days, FTW-mTSC teratomas were dissected and fixed with PBS containing 4% formaldehyde. For co-culture experiments, the FTW-mESC only group was injected with 1 × 10^6^ FTW-mESCs, and the co-culture group was injected with 1 × 10^6^ FTW-mESCs and 2.5 × 10^5^ FTW-mXENs. After one month, teratomas were dissected, weighed, and then fixed. Paraffin-embedded teratomas were sliced and stained with hematoxylin and eosin. Teratomas were formed using the following cell numbers: 1 × 10^6^ FTW-mESCs only, or 1 × 10^6^ FTW-mESCs plus 1.25 × 10^5^ (8:1), 2.5 × 10^5^ (4:1), 5 × 10^5^ (2:1), 1 × 10^6^ (1:1), 2 × 10^6^ (1:2) FTW-mXENs. After one month teratomas were dissected, weighed, and fixed.

### Immunostaining

Samples were fixed in 4% PFA for 15 min, washed three times with PBS and permeabilized with 0.5% Triton X-100 in PBS for 30 min at room temperature. Cells were then blocked with blocking buffer [5% (w/v) BSA; and 0.1% (v/v) Tween 20 in PBS] for 1 h and incubated with the primary antibodies (Table S7) diluted in blocking buffer at room temperature for 2 h or at 4°C overnight. After three washes with PBST (PBS plus 0.1% Tween 20), the cells were incubated with corresponding secondary antibodies (1:300 diluted, Table S7) in blocking buffer at room temperature for 1 h. After additional three times PBST washes, cells were counterstained with 300 nM DAPI solution at room temperature for 20 min before mounting. Samples were imaged using a fluorescence (Echo Laboratories, CA) or a confocal microscope (A1R, Nikon).

### Flow Cytometry

Cells were dissociated with TrypLE Express at 37 °C for 5 min. Then, the cells were fixed in 4% PFA at room temperature for 30 min and permeabilized with 0.5% (v/v) Triton X 100 at room temperature for 30 min. The cells were then incubated in the primary antibody (Table S7) solution for 30 min and then the secondary antibody solution for 30 min at room temperature. Samples stained with only secondary antibodies were used as the negative controls. Between each step, the samples were washed twice with PBS containing 2% FBS. Finally, the stained cells were suspended in PBS containing 2% FBS and analyzed by flow cytometry (FACScalibur system, BD).

### Blastocyst injection of FTW-mXENs and mouse FTW-mTSCs

FTW-mXEN injection into mouse blastocysts was performed as described previously(Yu *et al*., 2021b) with slight modifications. Briefly, single cell suspensions of mouse and human FTW-mXENs were added to a 40 μL droplet of KSOM-HEPES containing the blastocysts and placed on an inverted microscope (Nikon) fitted with micromanipulators (Narishige). Individual cells were collected into a micropipette with 15–20 μm internal diameter (ID), and a Piezo Micro Manipulator (Prime Tech) was used to create a hole in the zona pellucida and trophectoderm layer of mouse blastocysts. 10-12 (FTW-mXEN/ FTW-mTSCs) cells were introduced into the blastocoel. After microinjection, the blastocysts were cultured in mKSOMaa. For mouse embryo transfer, 8–12 weeks old ICR female mice were used as surrogates and were mated with vasectomized ICR male mice to induce pseudopregnancy. Ketamine (30 mg/ml) / Xylazine (4 mg/mL) and Buprenorphine (1 mg/mL) were used in surgery for maintaining anesthesia and relieving pain. Injected blastocysts were transferred to the surrogate uterine at E2.5. 14–30 blastocysts were transferred within 20–30 min per surrogate.

### Immunostaining and imaging of chimeric embryos

At E7.5 or E11.5, surrogates were euthanized, and embryos were isolated. Embryos were dissected and checked for fluorescence using zeiss Axio Zoom.V16 fluorescence stereo zoom microscope equipped with a Plan-Neofluar Z 1.0x/0.25 (FWD 56 mm) objective and Axiocam 503 monochromatic camera. Embryos were fixed in 4% paraformaldehyde and incubated at 4°C for 30 min (E7.5 embryos) or overnight (E11.5 embryos). After overnight cryoprotection in 30% sucrose solution (Fisher), the embryos were embedded in Polyfreeze Tissue freezing medium (Polyscience, Inc) and frozen on dry ice. Sections (10 μm thick) of the different embryos were cut on a Leica cryostat (Leica CM1950). For immunostaining, 10 mM citrate buffer (0.05% Tween 20 based) was used for antigen retrieval. The primary antibodies used were summarized in Table S7. After washing with TBST three times, the cells were incubated with corresponding secondary antibodies in blocking buffer at room temperature for 1 hour. Samples were counterstained with 300 nM DAPI solution at room temperature for 20 min and washed with PBST at least three times. Finally, slides were imaged using a fluorescence microscope (Echo Laboratories, CA).

### Mouse cell-cell co-culture assay

FTW-mESCs-eGFP, FTW-mTSCs-WT, and FTW-mXENs-mKO/WT were seeded onto MEF-coated plates either cultured separately or mixed at different ratios for co-cultures. The seeding ratio and density were empirically tested and decided on the basis of cell growth rate. A starting number of FTW-mESCs-eGFP (1.5 × 10^4^ cells), FTW-mTSCs-WT (3 × 10^4^ cells, and FTW-mXENs-mKO (7.5 × 10^3^ cells) were decided on for most of the cell-cell co-culture assays. During co-culture experiments, cells were cultured in FTW medium on MEFs for 5 days and the following analyses were performed. For co-culture experiments with various different cell ratios, FTW-mESCs-eGFP were seeded at a fixed cell number (1.5 × 10^4^ cells per well) and with varying numbers of FTW-mTSCs-WT and/or FTW-mXENs-WT.

For 3D co-culture, AggreWell™800 was used to form Embryoid bodies (EBs). For separate culture experiments, FTW-mESCs-eGFP were seeded into an AggreWell™800 with 2.4 × 10^5^ cells per well (800 cells per microwell), and for co-culture experiments, 2.4 × 10^5^ cells FTW-mESCs-eGFP and 1.2 × 10^5^ FTW-mXENs-WT were added per well (400 cells per microwell). The next day, EBs were transferred into 6-well low-attachment plates for further culture. Media was changed every day for 4 days using FTW medium. For 3D co-culture experiments with different ratios, FTW-mESCs-eGFP were seeded at the fixed density of 2.4 × 10^5^ cells per well and with varying densities of FTW-mXENs-WT.

For the differentiation co-culture experiments, FTW-mESCs-eGFP and FTW-mXENs-mKO were seeded, and the medium was switched to differentiation medium containing DMEM/F12 supplemented with 10% fetal bovine serum (FBS) the next day. On day 5, the number of FTW-mESCs-eGFP cells was counted.

### Transwell culture assay

For transwell co-culture experiments, Millipore Transwell 0.4 μm PET hanging inserts (Millicell, MCH12H48) were used by placing them into 12-well plates. MEFs were coated on the top/ bottom well. For separate culture groups, FTW-mESCs-eGFP (5000 cells) were seeded on the bottom wells, but not the top insert. For co-culture groups, FTW-mESCs-eGFP (5000 cells) and FTW-mXENs-WT (2500 cells) were seeded on the bottom and top insert of the wells, respectively. Half of the medium was changed every day and cell numbers were counted on day 5.

### Mouse ECM protein inhibition assay

Mouse FTW-mESC-GFP cells were passaged using TrypLE and seeded in a well of 6-well plate at 1.5 × 10^4^ cells in FTW cyESC medium under 20% O_2_ and 5% CO_2_ at 37. After 4 hours, the ECM protein mixtures (Matrigel_L: 0.5% (v/v), Matrigel_H: 2% (v/v), Laminin_L: 30 μg/ml, Laminin_H: 120 μg/ml, Collagen_L:15 μg/ml, Collagen_H:60 μg/ml, Vitronectin_L: 5 μg/ml, Vitronectin_H: 30 μg/ml) were directly added into the culture medium every day for 5 days. Medium and ECM proteins were changed every day.

### Monkey cell-cell co-culture assay

FTW-cyESCs-GFP, FTW-cyTSCs-WT, and FTW-cyXENs-mKO or WT were seeded onto MEF-coated plates and mixed at different ratios for co-culture experiments. The seeding ratios and densities were empirically tested and decided on the basis of cell growth rate. These numbers were: FTW-cyESCs-GFP (5 × 10^4^ cells), FTW-cyTSCs-WT (30 × 10^4^ cells, and FTW-cyXENs-mKO/WT (35 × 10^4^ cells). During co-culture experiments, cells were cultured in MEF-coated plates in FTW cyESC medium for 5 days.

### Monkey ECM protein inhibition assay

Monkey FTW-cyESC-GFP cells were passaged by treatment with TrypLE and 5 × 10^4^ cells were seeded into a well of 6-well plate in FTW medium under 20% O_2_ and 5% CO_2_ at 37. After 4 hours, ECM mixtures were added (Matrigel: 0.5%, 2%; Laminin: 2.5μg/ml, 10μg/ml; Collagen :15 μg/ml, 60 μg/ml) directly added into the culture medium every day for 5 days. Medium and ECM proteins were changed every day.

### Total FTW-mESCs/cyESCs number counting

For all of the co-culture experiments, total ES cell numbers were quantified in the following manner. Cells were dissociated into single cells using TrypLE Express at 37 °C for 4 min, and the total number of live cells were counted. Next, the relative percentage of eGFP+ FTW-ESCs-eGFP were determined using an LSR II Flow Cytometer (BD Bioscience). Total ES cell number (tN) for each group in both the co-cultured and separate culture conditions were determined by multiplying total cell volume (V) with cell concentration (CC) and percentage of eGFP+ cells (P). tN = V × CC × P. Cell density (cells cm^−2^) was calculated by dividing the total cell number by the surface area.

### Colony Size and intensity analysis

All of the co-culture experiments involving colony size and intensity were analyzed in the following method. On day 5 of co-culture, fluorescently labeled cells were imaged with a Leica Microsystem DMi8 microscope using Leica Application Suite X software for analysis. The images were taken randomly by the software in a fixed area for each well and A Fiji pipeline was used to quantify the size and intensity of the ESCs colonies. Briefly, images are preprocessed with median filters to filter out noise and small debris, binary images are then created through Otsu thresholding, and watershed was applied to separate merged clones. The quantification was implemented with the Analyze Particles function of Fiji based on the binary images created earlier. The average ESC colony Pixels (P) for each group in co-cultures and separate cultures were determined by multiplying colony Size (A) with colony intensity (I). P = A × I. For mouse cell experiments, there were three independent biological replicates for each group and 3 images for each sample. All of the colonies from 9 images were analyzed. For monkey cell experiments, 10 images were taken randomly and all of the colonies were analyzed in the 10 images.

### E6.5 epiblasts in-vitro cultivation assay with or without visceral endoderm

To obtain embryos, ICR females were mated with males from ICR (Charles River Laboratories) in the afternoon, and the presence of vaginal plugs was checked the next morning. The day on which a plug was found was considered to be E0.5. Both male and female mice were used at ages between 6 to 25 weeks. All the animal experiments were performed under the ethical guidelines of the Kindai University, and animal protocols were reviewed and approved by the Kindai University Animal Care and Use Committee. The developmental stage of embryos is critical for adapting the cultivation of isolated epiblast to an STO-conditioned medium. The isolated epiblasts before the onset of gastrulation collapsed in culture, while those after at onset of gastrulation were stable for growth in an STO-conditioned medium and were applicable for further experiments. The embryos at E6.5-6.75 were surgically isolated in cold DMEM supplemented with 10% fetal calf serum (FCS, BioWest), and whole epiblasts were isolated by the mechanical removal of Reichert’s membrane, extra-embryonic ectoderm as well as visceral endoderm, depending on the cases, using fine forceps and a tungsten needle. The isolated epiblasts with or without visceral endoderm were plated into Nunclon Sphera-treated, 96-well U-shaped-bottom microplate (174925, Thermo Scientific) in STO-conditioned medium. The STO-conditioned medium was prepared as follows. Dulbecco’s Modified Eagle’s Medium (DMEM, SIGMA) supplemented with 10% fetal bovine serum (FBS, Gibco) and 1% Penicillin-Streptomycin (10,000 U/mL, Gibco) was used to culture Mitomycin-C inactivated STO feeders for 2 days. After 48 hours, cultured epiblast outgrowths including visceral endoderm in the case were dissociated with TrypLE (Gibco, 12604013), and the total cell number was counted.

### Bulk RNA-sequencing

RNA extraction was performed using an RNeasy Mini Kit (QIAGEN) using DNase treatment (QIAGEN). RNA was analyzed using a 2100 Bioanalyzer (Aglient Technologies). (Transcripts per Kilobase Million). For monkey cells, RNA was extracted with Trizol Reagent (15596026, Invitrogen). The RNA (~50 ng) reverse transcription reaction and amplification were performed using SuperScript II (18064-071, Invitrogen), and KAPA HiFi HotStart Ready Mix (KK2602, KAPA). The cDNA was analyzed using a 2100 Bioanalyzer (Aglient Technologies). The RNA Library was generated using TruePrep DNA Library Prep Kit V2 for Illumina (TD501, Vazyme), then the library was adapted for sequencing on an Illmina NovaSeq 6000 platform (sequenced by Annoroad).

### Pre-processing of raw RNA-seq data

All reads were mapped to the mouse (GRCm38/mm10), human (GRCh38/hg38), and rhesus macaque genome (Mmul_10/rheMac10) using hisat2 (version 2.2.1) with default settings. FeatureCoumt (version 2.0.1) was used to estimate read counts. Stringtie (version 2.1.4) was used to estimate fragments per kilobase of exon per million fragments mapped (FPKM) and transcripts per kilobase of exon model per million mapped reads (TPM) values according to a previous report(Kovaka et al., 2019), genes with an FPKM value ≥3 were considered as expressed.

### Comparison analysis with published available datasets

The previously published datasets including mouse(Bao *et al*., 2018; Cruz-Molina *et al*., 2017; Cui *et al*., 2019; Kubaczka *et al*., 2015; Mohammed *et al*., 2017; Wu *et al*., 2015; Wu *et al*., 2011; Ye *et al*., 2018; Zhao *et al*., 2015; Zhong *et al*., 2018), human(Linneberg-Agerholm et al., 2019; Xiang et al., 2020; Zhou et al., 2019) and monkey(Ma *et al*., 2019; Nakamura *et al*., 2016; Niu *et al*., 2019b) embryonic single-cell RNA sequencing and mouse cell lines (nEND) microarray datasets(Anderson *et al*., 2017) were obtained from GEO repository (NCBI) and incorporated into our analysis. The expression levels of scRNA-seq data were transformed into log2(TPM + 1), and those of microarray data were transformed into log2(intensity).

### Principal components analysis (PCA)

The principal component analysis (PCA) was performed using the prcomp function without scaling. The DEGs were defined as genes exhibiting more than twofold changes between the samples (P < 0.005) and the sum of the expression level of every gene was log2(TPM+1) > 0 with the variance > 0.

### Similarities inference between cells

We selected the union of the top 2000 genes of highest variance for our dataset and the published dataset and calculated 20 canonical correlates (CCs) with diagonal CCA. After running CCA, the first 10 CCs were used for t-SNE visualization. The homology cell types were co-clustered in the same CCA cluster. Then, the correlation analysis was employed to detect the correspondence of cell subtype for our cells and the previous published embryonic cells by using expression matrix of 200 high variable genes that contributed to the first 10 CCs.

### Differential expression and GO and KEGG pathway analysis

Differentially expressed genes (DEGs) among clusters were detected by the Seurat function “Find All Markers”. Heatmaps showing the expression distribution of marker genes of cell clusters were created by pheatmap R package. We used the functions enrichKEGG and enrichGO in clusterProfiler R package(Yu et al., 2012) to perform KEGG pathways(Kanehisa et al., 2019) and Gene Ontology (GO) biological processes enrichment analysis. A pathway or process with a P value ≤ 0.05 was considered to be significantly enriched. The enriched pathways and processes were visualized with the ‘‘ggplot’’ function in the ggplot2 package in R.

### Single cell samples preparation

Mouse blastocysts were collected at E3.5 and the zona pellucida removed by Tyler’s buffer as described previously(Yu *et al*., 2021b). After removing the zona, blastocysts were washed in PBS/PVA two times. Digestion was performed in TrypLE Express for 10 minutes at 37, and cell clumps were cut up by mouth glass pipettes beginning with a larger size (50 um diameter) to a smaller size (20 um diameter). The totally digestion time was around 40-50 minutes. For the FTW-derive-day 8 outgrowth samples, blastocysts were collected and plated onto MEF-coated plates in FTW medium for 8 days, and then the outgrowth area were picked and dissociated. For the established FTW cell lines, all the cells were dissociated with TripLE Express for 3-5 minutes at 37.

### scRNA-seq library preparing and sequencing

Single cell suspensions were diluted at a concentration of 1100 cells/µL in 0.04% bovine serum albumin (BSA)/PBS for loading into 10X Chromium Single Cell G Chips (for mouse blastocysts, all the of the cells were loaded into the chip). Single cell libraries were prepared using the Chromium Single Cell 3’ Reagent Kit v3.1 (10X Genomics, Pleasanton, CA) following the manufacturer’s protocol. Briefly, single cells were partitioned into Gel beads in EMulsion (GEMs) in the GemCode instrument followed by cell lysis and barcoded reverse transcription of RNA, amplification, shearing and 5’ adaptor and sample index attachment. Libraries were sequenced on the Illumina NovaSeq 6000 platform.

### scRNA-seq data analysis

Single-cell gene expression count matrix was constructed with Cell Ranger (v6.0.0). The subsequent analyses were performed using the Seurat package (v4.6.0) (Hao et al., 2021). For each dataset, transcription noise cells were removed by a series of criteria, including minimal expression of 200 genes per cell, mitochondrial read percentage < 20%, ribosomal read percentage > 5%, as well as specific minimal/maximal number of genes and maximal reads for each dataset. For the monkey, after stringent quality filtering, we generated 35987 single cell transcriptomes with a median unique molecular identifier (UMI) of 22277 and gene number of 5436. Cells passed quality control were merged and normalized using Seurat’s SCT method. Datasets from different sources were integrated using Seurat’s CCA method. The UMAP dimensional reduction was performed using the first 30 PCs from the PCA analysis. Cells were then clustered using the FindClusters function in Seurat package. Cell identities were annotated based on the expression of cell marker genes. To examine the correlation among cell clusters, we first performed PCA analysis for each cell cluster and then carried out correlation analysis using the PC1s extracted from the cell clusters.

### Cell clustering by nonlinear dimensional reduction

To remove the influence of different experimental conditions on the samples, batch effect correction was carried out. Briefly, following the standard processing steps, including use of “SelectIntegrationFeatures” function to select 2,000 feature genes, “FindIntegrationAnchors” function to find a set of anchors between different samples, “IntegrateData” function to integrate data, “RunPCA” and “RunUMAP” function select 30 PCs dimensionality reduction, “DimPlot” function to graphs the output of a dimensional reduction technique on a 2D scatter plot. The Seurat package was applied to perform cell clustering analysis based on TPM expression values, and only the genes with an expression level greater than 5 were included for further analysis. Then uniform manifold approximation and projection (UMAP) were applied for visualization. First, ‘‘FindIntegrationAnchors’’ function took these objects as input with parameters as ‘‘k.anchor = 5, anchor.features = 2000’’ and returned an anchor object. Then, ‘‘IntegrateData’’ function used the anchor object to integrate all data-sets with the default parameter. Finally, the top-10/30 principal components (PCs) were used for clustering (with a resolution of 1) by Seurat.

### Pseudotime trajectory analysis

Slingshot(Street *et al*., 2018) was used to determine cell Pseudotime trajectories. Frist, the object that determines the cell type is passed into the “SingleCellExperiment” sim object. Then use a “Rtsne” function to dimensionality reduction (dims = 2, perplexity = 10, pca = TURE, pca_scale = FALSE, normalize = FALSE). Slingshot analysis was then performed on the diffusion map to determine per-cell pseudotime estimates and mapped back to the UMAP embedding.

### Gene regulatory networks

Single cell regulatory network inference and clustering (SCENIC) (Aibar et al., 2017) has been used to infer gene regulatory networks based on single cell expression profiles and identify cell states, providing important biological insights into the mechanism of cell heterogeneity. To identify important transcriptional regulation during cell development, we used pySCENIC grn/ctx/aucell of the Python module tool pySCENIC to obtain core regulatory TF. The workflow first describes the input single-cell expression abundance profile matrix and applies a per-target regression method (GRNBoost2) to infer co-expression modules from which indirect targets are pruned based on cis-regulatory motif discovery (cisTarget). Subsequently, the activity of these regulatory factors was quantified using aucells, and the regulatory factor activity score (RAS) was obtained by enrichment and scoring the target genes of the regulatory factors. The regulator specificity score (RSS) was calculated based on Jensen-Shannon divergence (JS) to determine the cluster-specific regulators, this process is implemented through R language “calcRSS” function and visualized through ggplot2.

### Cell-cell communication analysis

To compare the changes of the cell-cell interactions among ES, TS and XEN between the co-culture and separate culture conditions, we performed cell-cell communication analysis using the CellChat package(Jin *et al*., 2021). Following the official workflow, we calculated the potential ligand-receptor interactions among the three cell lineages in each condition and then compared the numbers and strength of these interactions between the two conditions. Briefly, loaded the normalized counts into CellChat and applied the standard preprocessing steps, including the functions “identifyOverExpressedGenes”, “identifyOverExpressedInteractions”, and “projectData” with a standard parameter set. We then calculated the potential ligand-receptor interactions between infected and non-infected cells based on the functions “computeCommunProb”, “computeCommunProbPathway”, and “aggregateNet” using standard parameters, all of the above procedures use default parameters.

### CRISPR knockout

We used the online software Benchling CRISPR to design all single guide RNAs (sgRNAs) used in this study. The sequences of sgRNAs are included in Table S7. sgRNAs were cloned into the pSpCas9(BB)-2A-eGFP (Addgene, PX458) plasmid by ligating annealed oligonucleotides with BbsI-digested vector. The plasmid carrying the specific sgRNA was then transfected into FTW-mESCs or FTW-mXENs by using an electroporator (NEPA2, Nepa Gene 1). EGFP positive cells were sorted by flow cytometry after 48 h transfection and 2000 cells were plated in one well of a 6 well plate. Single clones were picked and expanded. Homozygous knockout clones were confirmed by Sanger sequencing.

### Plasmids

pSpCas9(BB)-2A-eGFP (PX458) plasmid was purchased from Addgene (plasmid 48138). pCAG-IP-mKO, pCAG-IP-eGFP plasmids were obtained from T. Hishida.

### Statistical Analysis

All quantitative data is presented as the mean ± SD. Experiments were repeated at least three times (repeat number was indicated as “n” in figure legends). Differences between groups were evaluated by Student’s t-test (two-sided). *P* values are shown in the figures. Graphic analyses were done using GraphPad Prism version 7.0 and 8.0 (GraphPad Software, La Jolla, Ca) and Microsoft Excel (Microsoft 365).

---

**Figure S1.**
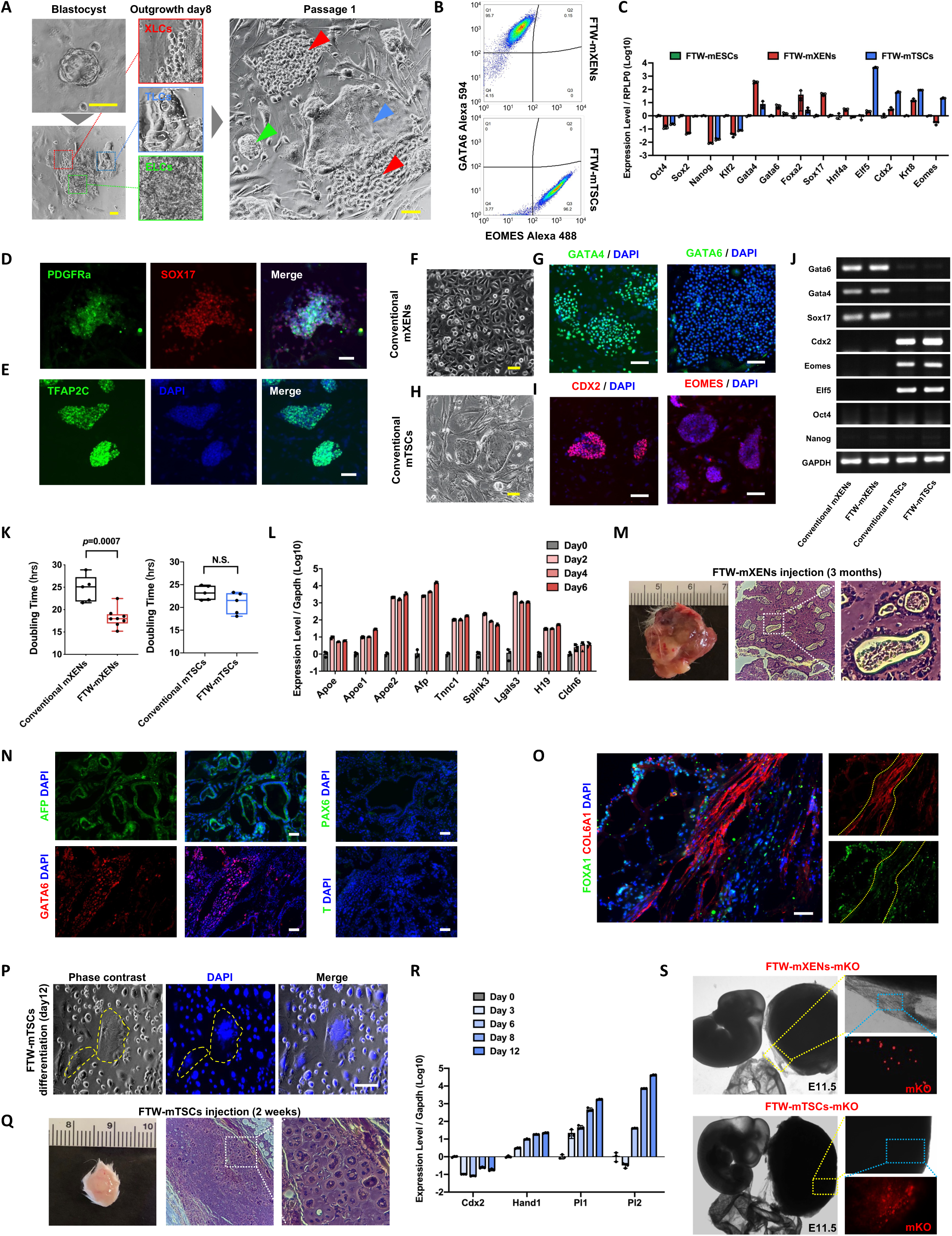
Derivation, characterization of FTW-mXENs, FTW-mTSCs and FTW-mESCs and developmental potential of FTW-mXENs and FTW-mTSCs. (A) Representative BF images of an E4.5 mouse blastocyst (upper left), day 8 outgrowth (bottom left) and passage 1 (right) during FTW stem cells derivation. Higher magnification images of the boxed areas are shown in the middle panel. Scale bars, 100 µm. (B) Flow cytometry results showing the expression of GATA6 and EOMES in FTW-mXENs (left) and FTW-mTSCs (right). (C) qRT-PCR results showing the relative expression levels of lineage markers in FTW-mESCs, FTW-mXENs and FTW-mTSCs. (mean ± SD, n = 3, biological replicates). (D) Representative IF images of PDGFRa and SOX17 of FTW-mXENs. Scale bar, 100 µm. (E) A representative IF image of TFAP2c of FTW-mTSCs. Scale bar, 100 µm. (F) to (I), Representative BF (F and H) and IF images (G and I) of conventional mXENs (F and G) and mTSCs (H and I). Scale bars, 100 µm. (J) RT-PCR results showing the expression of lineage markers in conventional mXENs and mTSCs and converted FTW-mXENs and FTW-mTSCs. GAPDH, loading control. (K) Doubling time of conventional mXENs (n = 5, biological replicates), mTSCs (n = 5, biological replicates), converted FTW-mXENs (n = 8, biological replicates), and converted FTW-mTSCs (n = 5, biological replicates). Box-and-whisker plots showing the median value (bar inside box), 25^th^ and 75^th^ percentiles (bottom and top of box, respectively), and minimum and maximum values (bottom and top whisker, respectively). (L) qRT-PCR results showing the relative expression levels of several PE and VE markers in randomly differentiated FTW-mXENs at indicated timepoints (mean ± SD, n = 3, biological replicates). (M) Representative images showing the appearance and H&E staining of a XEN-teratoma. A higher magnification image of the boxed area is shown on the right. (N) Representative IF images of AFP, GATA6, PAX6 and T in XEN-teratoma sections. Blue, DAPI. Scale bars, 100 µm. (O) A representative IF image of FOXA1 and COL6A1in a XEN-teratoma section. (P) Representative BF and DAPI staining image of randomly differentiated FTW-mTSCs at day 12. Yellow dashed lines indicate multinucleated cells. Scale bar, 100 µm. (Q) Representative images showing the appearance and H&E staining of a TSC-teratoma. A higher magnification image of the boxed area is shown in the right panel. (R) qRT-PCR results showing the relative expression levels of trophoblast differentiation-associated genes in random differentiated FTW-mTSCs at indicated timepoints (mean ± SD, n = 3, biological replicates). (S) Representative BF and fluorescence images showing chimeric contribution of mKO-labeled FTW-mXENs (left) and FTW-mTSCs (right) to E11.5 mouse conceptuses. Scale bars, 1 mm.

**Figure S2.**
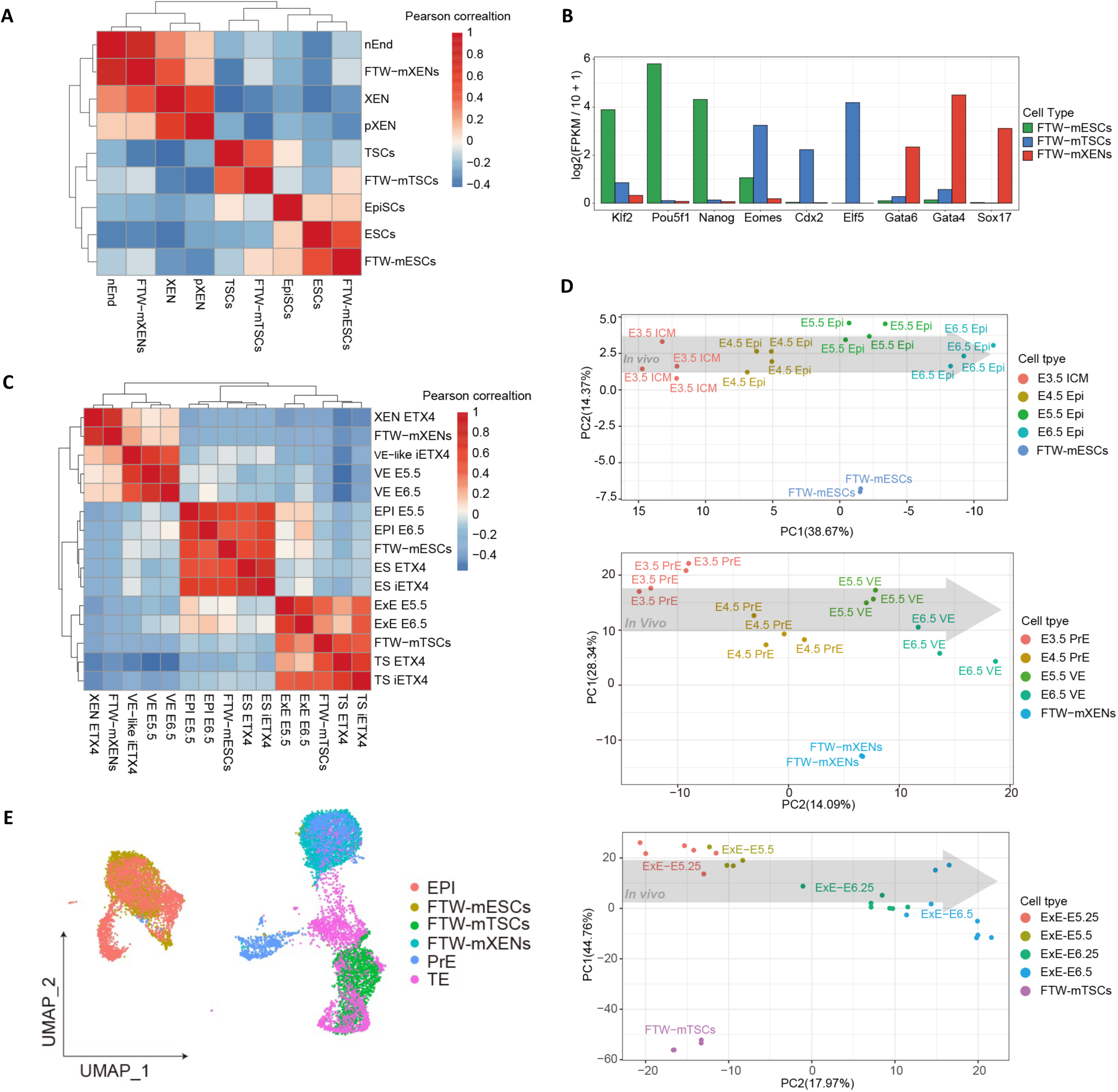
Transcriptomic analyses of mouse FTW stem cells. (A) Correlation analysis of FTW-mXENs, FTW-mTSCs, FTW-mESCs and published datasets of mPSCs(Bao et al., 2018; Cruz-Molina et al., 2017; Wu et al., 2015; Ye et al., 2018; Zhao et al., 2015), mTSCs(Cui et al., 2019; Kubaczka et al., 2015; Wu et al., 2011) and mXENs(Anderson et al., 2017; Zhong et al., 2018). (B) The expression levels (log2 [FPKM\10+1]) of epiblast, trophoblast and extraembryonic endoderm markers in FTW-mESCs, FTW-mTSCs, and FTW-mXENs obtained from RNA-seq datasets. (C) Correlation analysis of FTW-mXENs, FTW-mTSCs, FTW-mESCs and published datasets from mouse embryos and stem cell embryo models. (D) PCA plots of RNA-seq datasets from FTW-mESCs, FTW-mTSCs, and FTW-mXENs and in vivo E3.5-E6.5 mouse conceptuses. Inner cell mass (ICM), Epiblast (Epi), Primitive endoderm (PrE), Visceral endoderm (VE), Extraembryonic endoderm (ExE). (E) UMAP analysis of scRNA-seq datasets of FTW-mESCs, FTW-mTSCs, FTW-mXENs and in vivo E5.5 mouse conceptuses (EPI, PrE and TE cells).

**Figure S3.**
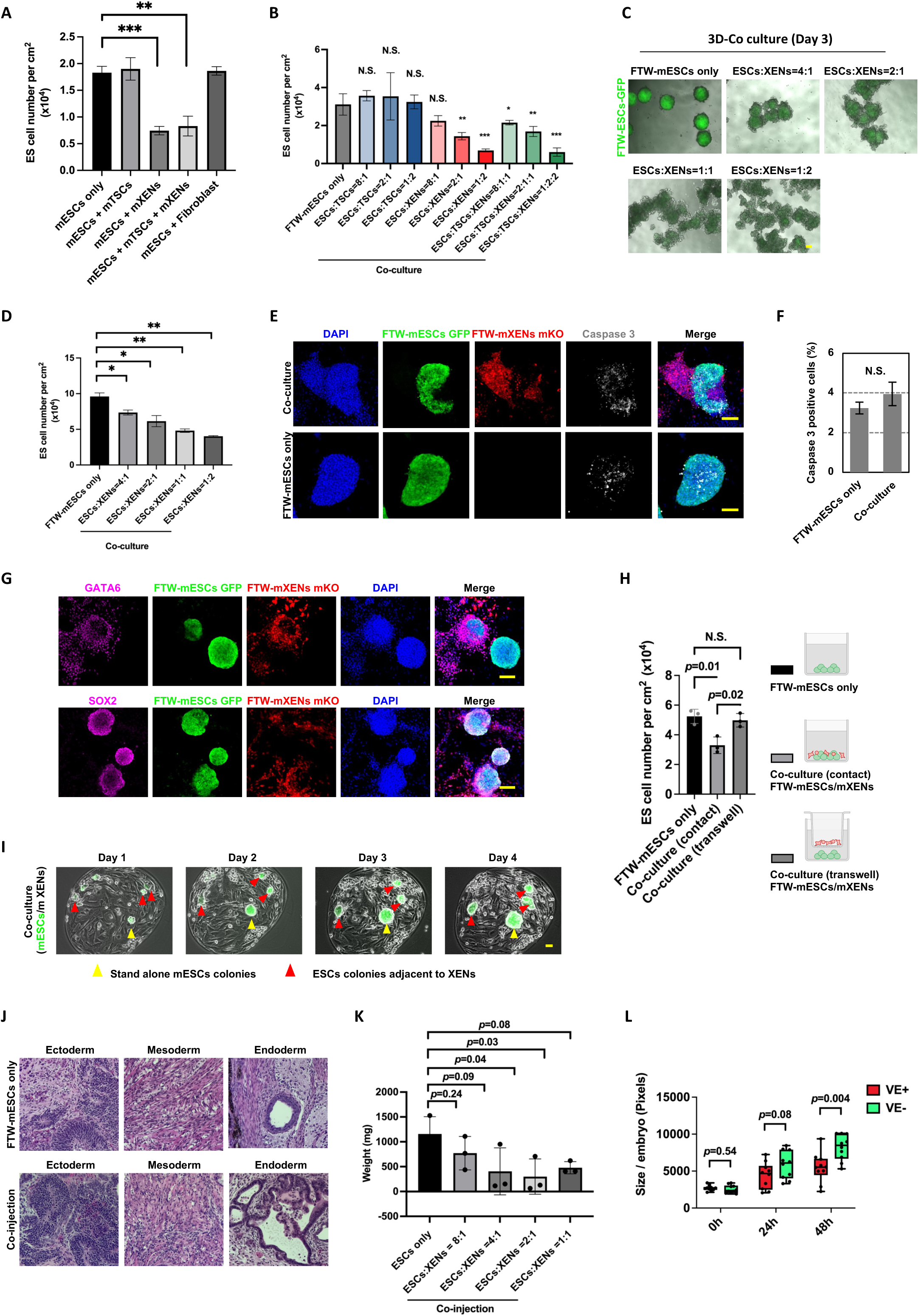
Proliferation restriction of FTW-mESCs by FTW-mXENs. (A) Bar plot showing the cell densities of FTW-mESCs in day 5 separate and different co-cultures. (B) Bar plot showing the cell densities of FTW-mESCs in separate and co-cultures (different mESCs: mTSCs and/or mESCs:mXENs ratios) (C) Representative BF and fluorescence merged images of day 5 3D separate and co-cultures (different mESCs:mXENs ratios). Green, FTW-mESCs. Scale bar, 100 µm. (D) Bar plot showing the total cell number of day 5 3D separate and co-cultures (different mESCs:mXENs ratios) (mean ± SD, n = 2, biological replicates). (E) Representative IF images showing the expression of Activated Caspase 3 in separately cultured FTW-mESCs (bottom) and FTW-mESCs co-cultured with FTW-mXENs (top). Scale bars, 100 µm. (F) Bar plot showing the percentages of Caspase3+ cells in separately cultured FTW-mESCs (left) and FTW-ESCs co-cultured with FTW-mXENs (right). (mean ± SD, n = 3, biological replicates, N.S., not significant). (G) Representative IF images showing the expression of GATA6 and SOX2 in co-cultured FTW-mXENs (mKO) and FTW-mESCs (GFP), respectively. Scale bars, 100 µm. (H) Bar plot showing the cell densities of FTW-mESCs in day 5 separate, contact and non-contact (transwell) co-cultures (mean ± SD, n = 3, biological replicates). (I) Representative BF and fluorescence merged images of FTW-mESCs and FTW-mXENs in micropatterned co-cultures from day 1 to day 4. Scale bars, 100 µm. Yellow arrowhead, a standalone FTW-mESCs colony. Red arrowheads, FTW-mESCs colonies adjacent to FTW-mXENs. (J) Representative H&E staining images of a teratoma generated from FTW-mESCs alone (top) and FTW-mESCs co-injected with FTW-mXENs (bottom). (K) Weight of teratoma formed by FTW-mESCs and FTW-mESCs co-injected with FTW-mXENs at different mix ratios (mean ± SD, n = 3, biological replicates). (L) Bar plot showing the area of each in vitro cultured E6.5-6.75 mouse epiblast with (red) or without (green) VE (mean ± SD, n = 10, biological replicates). *P-value < 0.01, **P-value < 0.001. P-values were calculated using two-tailed Student’s t test.

**Figure S4.**
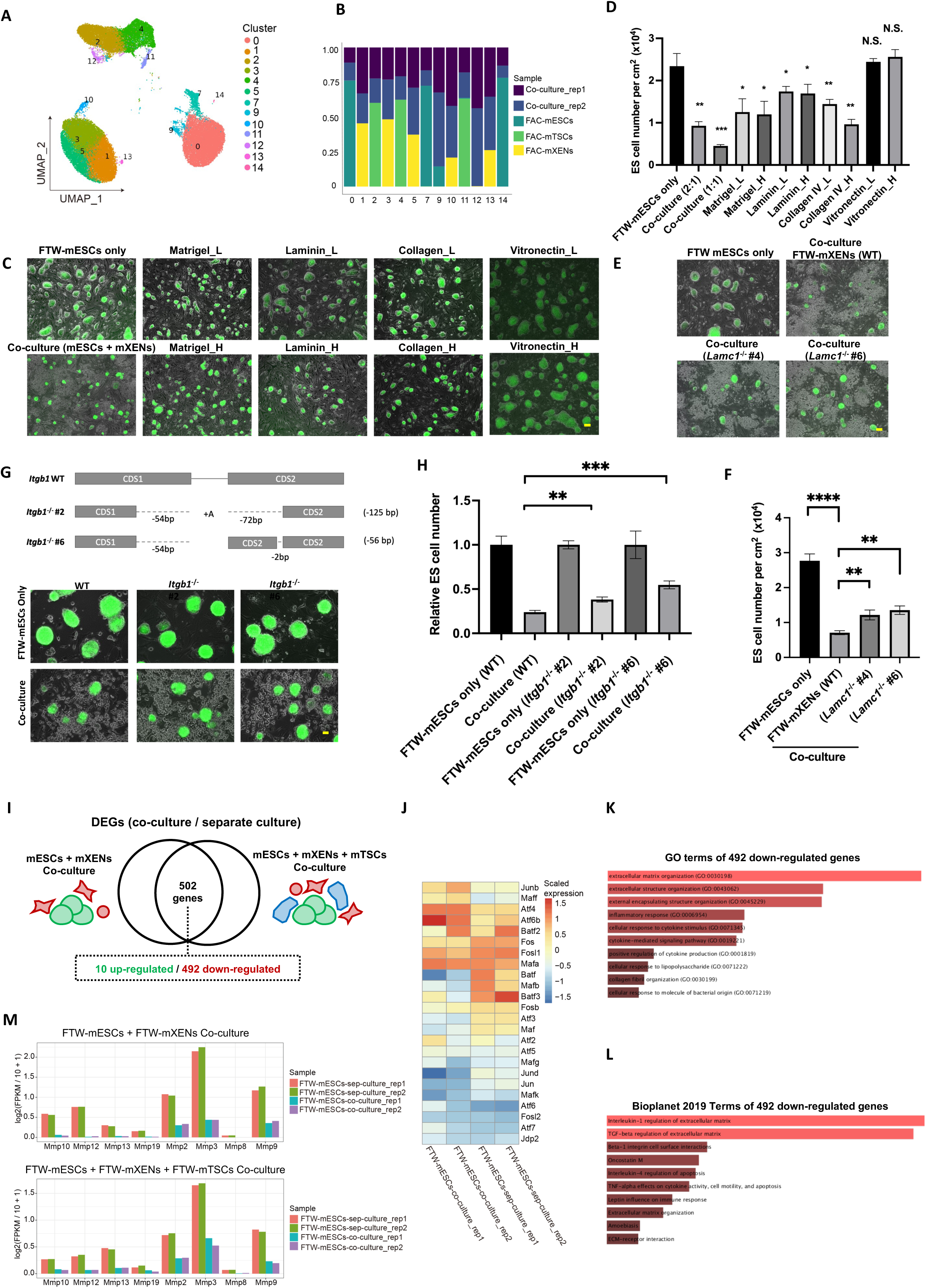
Mechanistic insights of growth inhibition and transcriptomic analyses of co-cultured FTW stem cells. (A) Joint UMAP embedding of single-cell transcriptomes from separately- and co-cultured FTW-mESCs, FTW-mXENs or FTW-mTSCs. (B) Percentage of cells in each cluster derived from separately and co-cultured FTW-mESCs, FTW-mXENs or FTW-mTSCs. (C) Representative BF and fluorescence merged images of separately- and co-cultured FTW-mESCs, as well as separately cultured FTW-mESCs supplemented with different ECM proteins. Scale bar, 100 µm. (D) Bar plot showing the cell densities of FTW-mESCs in day 5 separate and co-cultures, as well as separate cultures supplemented with different ECM proteins (mean ± SD, n = 3, biological replicates, N.S., not significant). (E) Representative BF and fluorescence merged images of separately- and co-cultured FTW-mESCs (with WT or *Lamc^−/−^* FTW-mXENs [clones #4 and #6]). (F) Bar plot showing the cell densities of FTW-mESCs in day 5 separate and co-cultures (with WT of *Laminin^−/−^* FTW-mXENs) (mean ± SD, n = 3, biological replicates). (G) Representative BF and fluorescence merged images of separately and co-cultured FTW-mESCs (WT or *Itgb1^−/−^* [clones #2 and #6]) Scale bar, 100 µm. (H) Bar plot showing the relative cell densities of FTW-mESCs (WT or *Integrin-β1^−/−^*) in day 5 separate and co-cultures (mean ± SD, n = 3, biological replicates). (I) A VENN diagram showing shared 502 DEGs (492 down-regulated; 10 up-regulated) between separately cultured FTW-mESCs and FTW-mESCs co-cultured with FTW-mXENs or with both FTW-mXENs and FTW-mTSCs. (J) A heatmap showing the expression levels of AP-1 family members in separately (FTW-mESCs) and co-cultured (FTW-mESCs-co) FTW-mESCs. (K) and (L), Enriched GO terms (K) and Bioplanet terms (L) for 492 down-regulated genes from (I). (M) MMP expression levels (FPKM value) in separately cultured FTW-mESCs, FTW-mESCs co-cultured with FTW-mXEN (top), and FTW-mESCs co-cultured with FTW-mXENs and FTW-mTSCs (bottom).

**Figure S5.**
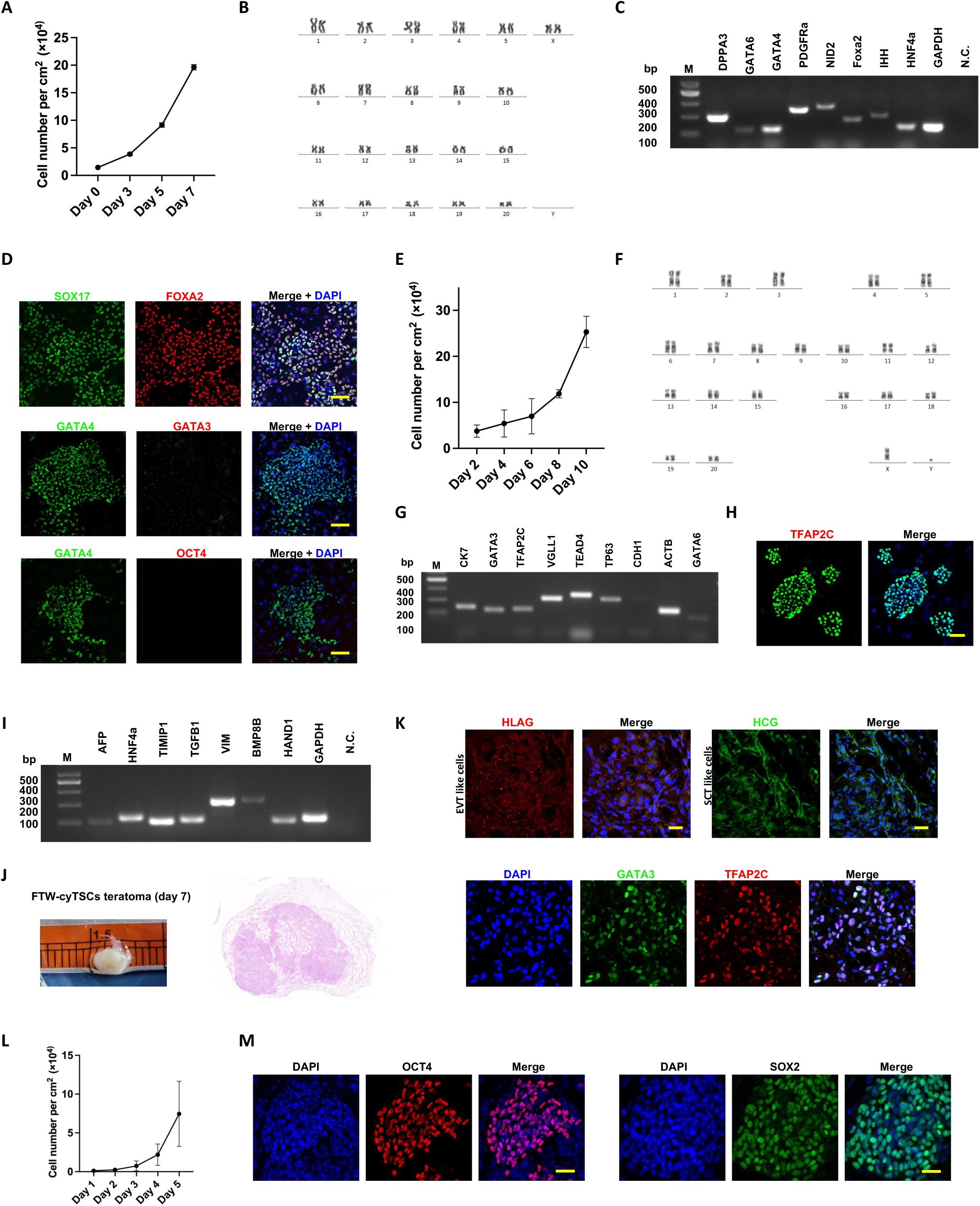
Derivation and characterization of FTW-cyXENs, FTW-cyTSCs and FTW-cyESCs. (A) Growth curve of FTW-cyXENs (passage 26) (n =9, biological replicates). (B) Karyotype analysis of FTW-cyXENs. (C) RT-PCR results showing the expression of several hypoblast markers in FTW-cyXENs. M, DNA ladder. N.C., non-template control. (D) Representative IF images showing GATA4, SOX17, FOXA2, GATA3 and OCT4 expression patterns in FTW-cyXENs. Blue, DAPI. Scale bars, 50 µm. (E) Growth curve analysis of FTW-cyTSCs (passage 16) (n = 2, biological replicates). (F) Karyotype analysis of FTW-cyTSCs. (G) RT-PCR results showing the expression of several trophoblast markers in FTW-cyTSCs. M, DNA ladder. N.C., non-template control. (H) Representative IF images showing TFAP2C expression in FTW-cyTSCs. Blue, DAPI. Scale bar, 50 µm. (I) RT-PCR results showing the expression of VE/YE and EXMC related genes in differentiated FTW-cyXENs at day 9. (J) Representative BF and H&E staining images of a TSC-teratoma generated from FTW-cyTSCs at day 7 post-injection. (K) Representative IF images showing the expression patterns of EVT (HLAG), SCT (HCG), and TSC (GATA3 and TFAP2C) marker genes in FTW-cyTSCs derived TSC-teratoma sections. Blue, DAPI. Scale bars, 50 µm. (L) Growth curve of FTW-cyESCs (passage 9) (n = 3, biological replicates). (M) Representative IF images showing the expression of OCT4 and SOX2 in FTW-cyESCs. Blue, DAPI. Scale bars, 50 µm.

**Figure S6.**
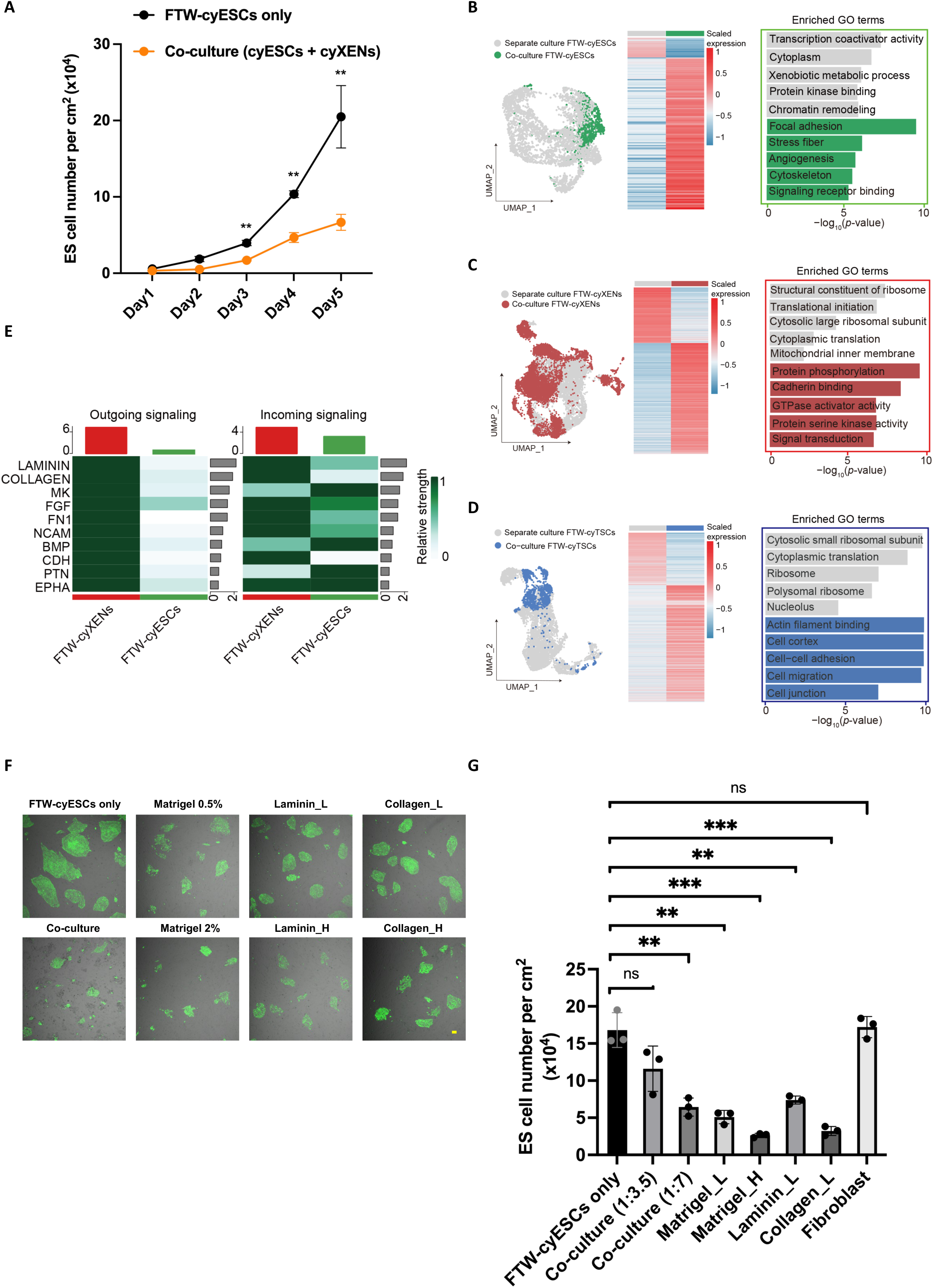
Lineage crosstalk among co-cultured cynomolgus monkey FTW stem cells. (A) Growth curves of separately and co-cultured (with FTW-cyXENs) tFTW-cyESCs (n=3, biological replicates). (B) to (D), DEGs of separately cultured and co-cultured FTW-cyESCs (B), FTW-cyXENs (C), and FTW-cyTSCs (D). Top enriched GO terms were shown on the right. (E) Heatmaps showing top 10 outgoing (left) and incoming (right) signaling in co-culture of FTW-cyESCs and FTW-cyXENs. (F) Representative BF and fluorescence merged images of separately- and co-cultured FTW-cyESCs, as well as separately cultured FTW-cyESCs supplemented with different ECM proteins. Scale bar, 100 µm. (G) Bar plot showing the cell densities of FTW-cyESCs in day 5 separate and co-cultures, as well as separate cultures supplemented with different ECM proteins (n=3, biological replicates).

**Figure S7.**
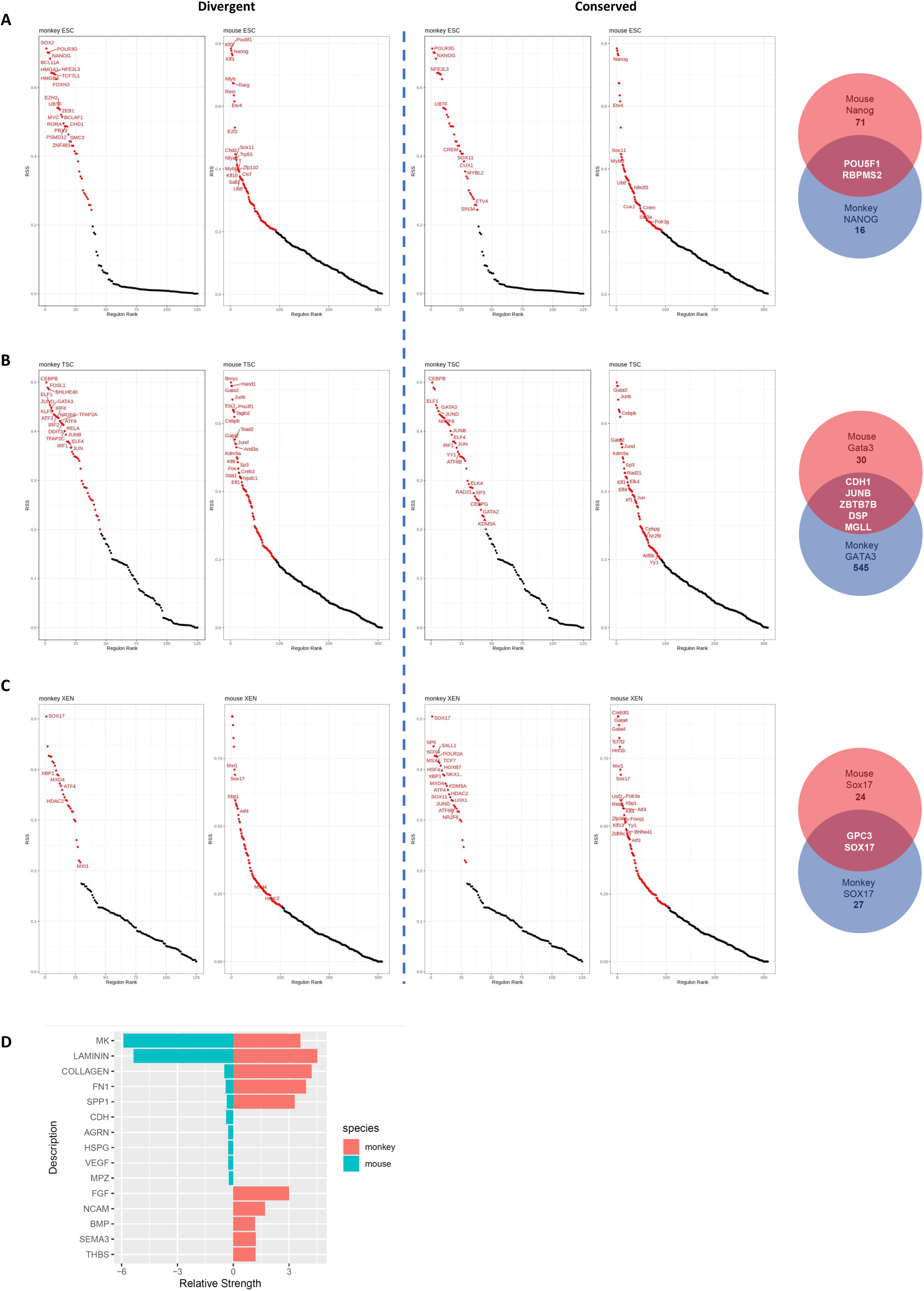
Cross-species comparison. (A) to (C) SCENIC analysis showing divergent (left) and conserved (middle) TF-driven regulons in each mouse and monkey stem cell. Right, VENN plots showing common and distinct targets (between mice and cynomolgus monkeys) of well-known lineage TFs including NANOG, SOX17 and GATA3. See Table S6 for the all the target lists. Red dots represent the gene regulatory networks regulated by corresponding transcription factor related to the RBPS. (D) Cross-species comparison of signaling crosstalk among embryonic and extraembryonic cells.

## Notes

### Summary of Updates

Order or the authors appear on the bioRxiv website.

